# Metabolic Plasticity drives Development during Mammalian Embryogenesis

**DOI:** 10.1101/2020.10.07.330571

**Authors:** Mark S. Sharpley, Fangtao Chi, Utpal Banerjee

## Abstract

Preimplantation mouse embryos interact minimally with their environment, and development is largely driven by metabolic processes. During the earliest cleavage stages, metabolism is rigid, with maternal deposits enforcing a redox state that facilitates zygotic genome activation. As maternal control falls, metabolic shuttles are activated, increasing glycolysis and equilibrating the TCA cycle. The resulting flexibility of nutrient utilization and metabolic plasticity facilitates unidirectional developmental progression such that later stage embryos proceed to form blastocysts without any exogenously added nutrients. We explore the mechanisms that govern this choreographed sequence that balances the deposition, degradation, synthesis and function of metabolic enzymes with redox control, bioenergetics and biosynthesis. Cancer cells follow a distinct metabolic strategy from that of the preimplantation embryo. However, important shared features emerge under reductive stress. We conclude that metabolic plasticity drives normal development while stress conditions mimic hallmark events in Cancer Metabolism.

## Introduction

During the course of mammalian preimplantation development, a single-cell, the zygote, develops into the blastocyst, which is a complex multicellular structure (Rossant, 2018). In mice, the formation of the blastocyst takes 4-5 days. Zygotic genome activation (ZGA) occurs at the two-cell (2C) stage of development, and prior to this time the embryo largely relies on proteins or mRNAs that are inherited from the oocyte. At the 8-cell stage, the mouse embryo undergoes compaction, during which an embryo, that initially comprises loosely adhered cells, morphologically transforms into a tightly connected mass of cells giving rise to a compacted morula (White et al., 2016). Following this morula stage transition, the outer cells of the embryo differentiate to form trophectoderm (TE) precursors that will contribute to the placenta, whereas the inner cells will form the inner cell mass (ICM) that will contribute to all embryonic and some extraembryonic (primitive endoderm, PE) tissue (Leung et al., 2016). Following blastocyst formation, the ICM will form the epiblast, from which pluripotent embryonic stem cells (ESCs) are derived.

The importance of metabolism in controlling preimplantation embryo development was established several decades ago by pioneers who identified conditions that enable early embryos to grow outside of the oviduct (Biggers et al., 1967; Brinster, 1963; Leese, 2012). A key insight gained during these early studies was that embryos have an obligate requirement for pyruvate to develop beyond the 1-2 cell stage (Figure 1A). From the late 2-cell stage, either pyruvate or lactate can facilitate development (Brown and Whittingham, 1991; Lane and Gardner, 2005). Glucose or glutamine (Gln), alone or in combination, cannot support the development of embryos at this stage, which contrasts with cultured cancer cells for which these metabolites are the major source of energy and biomass (Altman et al., 2016; Liberti and Locasale, 2016; Pavlova and Thompson, 2016; Vander Heiden and DeBerardinis, 2017).

**Figure 1.**
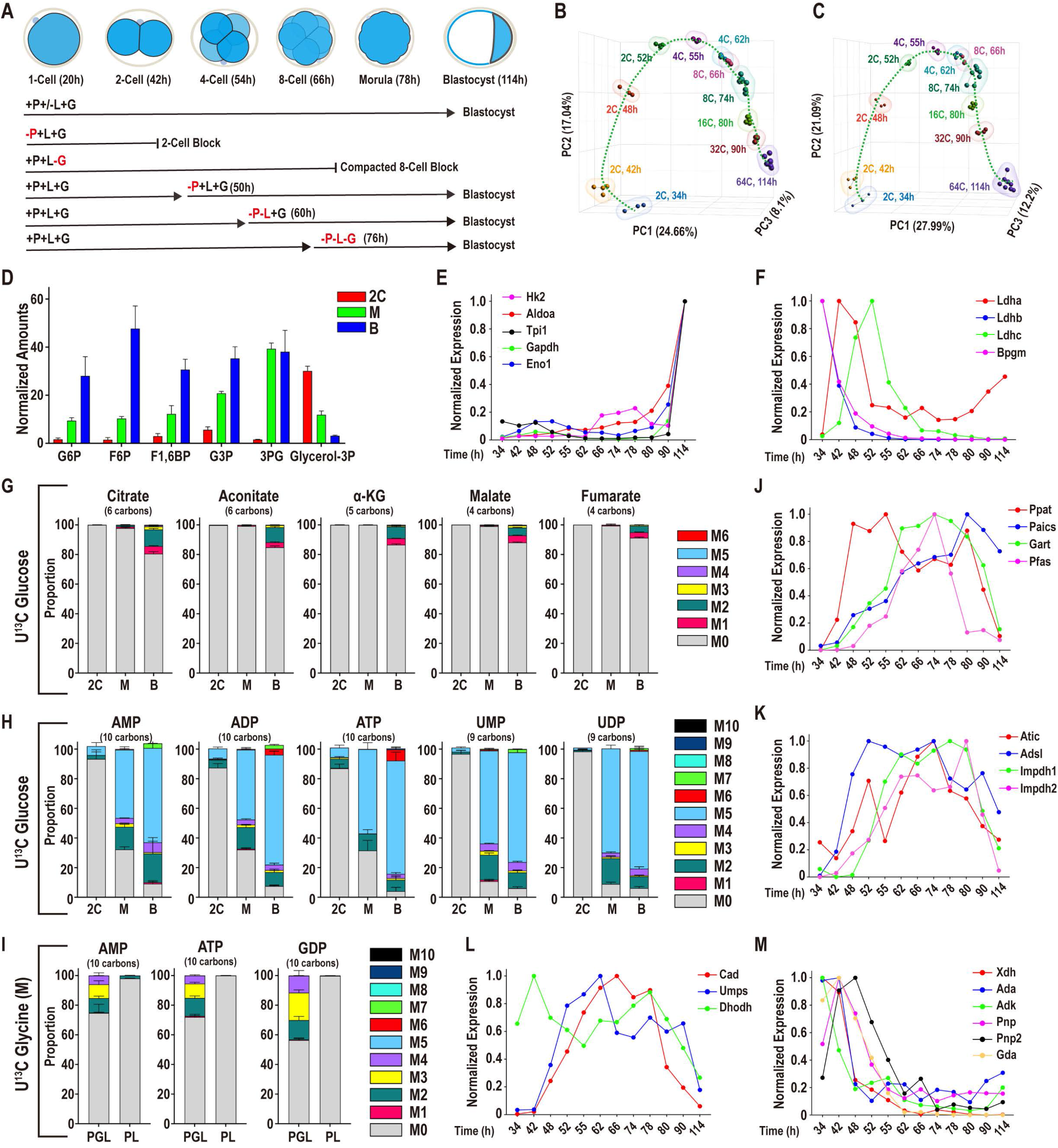
Glucose metabolism during preimplantation development. In this and in all figures, times in hours (h) refer to time elapsed after human chorionic gonadotropin (hCG) injection that induces ovulation. Zygotes are isolated at 18h and cultured until the specified hours (h) post hCG. The metabolomic analysis is presented as scaled means ± SD and is obtained from 3 biological replicates. **(A**) A schematic representation of developmental progression of mouse preimplantation embryos in the presence of different combinations of nutrients. Based on past literature and confirmed here, embryos that are cultured from the zygote stage with glucose and lactate, but without pyruvate, block at the 1C and 2C stages and embryos cultured without glucose block at the 8-16C (morula) stage. Following the late 2C stage, embryos cultured without pyruvate develop into blastocysts as long as lactate and glucose are present. Later in this paper we demonstrate that after 60h the embryo is able to develop with only glucose provided, and also that following 76h, no exogenously provided nutrients are required for blastocyst formation. **(B, C)** Principal component analysis (PCA) of changes in gene expression during preimplantation development. **(B)** PCA for the entire transcriptome (34112 genes), and **(C)** PCA limited to genes that are directly involved in metabolic processes (1947 genes). The characteristic inverted U-shaped clustering for both **(B)** and **(C)** indicates a dynamic control of metabolic genes during development. **(D)** Relative amounts of glycolytic intermediates in 2C (48h), morula (M, 78h), and blastocysts (B, 96h). For each metabolite, the bars provide indications of relative scaled amounts and therefore this chart is not intended for comparisons of the absolute amounts for different metabolites. **(E, F)** Expression of genes encoding glycolytic enzymes between early 2C (34h) and fully expanded blastocyst (114h) stages. **(E)** A majority of glycolytic enzymes increase in expression during later stages of development. **(F)** A subset is highest at the 2C stage and then declines. See also Figure S1. **(G)** U-^13^C glucose only contributes minor amounts of carbon to TCA cycle metabolites, with a measurable contribution only at the blastocyst stage. Gray: M0 unlabeled; colors: labeled as marked. Note that the “fully labeled” color depends on the total carbons in the metabolite (e.g. fully labeled is M6 for citrate, M5 for alpha-KG and M4 for fumarate). **(H)** U-^13^C glucose contributes to nucleotide formation in increasing amounts during development. The major isotopologue, M5 (blue) represents fully labeled ribose formation by glucose. **(I)** U-^13^C glycine contributes to purine base synthesis in morulae if glucose (U-^12^C) is present (PGL), but does not do so when glucose is withheld (PL). **(J, K, L, M)** Expression of nucleotide metabolism genes. A majority of the *de novo* purine (**J, K**) and pyrimidine (**L**) synthesis genes increase in expression during later stages, whereas genes involved in purine degradation (**M**) peak in expression during the 2C stage.

In spite of extensive earlier metabolic studies direct and comprehensive metabolomic measurements are lacking for the normal development of the preimplantation embryo, largely due to a paucity of available biological material (Barbehenn et al., 1974; Lane and Gardner, 2000; Leese and Barton, 1984; Shyh-Chang et al., 2013; Wales, 1975; Zhang et al., 2018). This contrasts with the extensive metabolomic analyses in the literature for cultured cancer cells and ESCs (Faubert et al., 2020; Intlekofer and Finley, 2019; Jang et al., 2018; Zhang et al., 2016).

In the past, we have shown that metabolism impacts several signaling processes that control preimplantation development (Chi et al., 2020; Nagaraj et al., 2017). For example, at the 2-cell stage, ZGA requires a novel pyruvate-mediated translocation of mitochondrial TCA cycle enzymes to the nucleus (Nagaraj et al., 2017), where these enzymes generate metabolites essential for the epigenetic changes and required for genome-level reprogramming of transcription. Later, at the morula stage, a glucose-mediated signaling process is critical for the control of expression of transcription factors necessary for TE, but not ICM, cell differentiation, and this distinction facilitates the first cell fate-choice of the embryo (Chi et al., 2020). In this manuscript, we present the first comprehensive stable isotope-resolved metabolomic analysis across all stages of preimplantation development and correlate these results with transcriptomic analysis to characterize the capacity of the embryo to reprogram its metabolism during normal development and in response to changes in the environment. In doing so we are able to elucidate the basis for metabolic plasticity as a driver of developmental progression of the mammalian embryo.

## Results

### Glucose metabolism during preimplantation development

At the morula (8-16 cell) stage, glycolysis is dispensable even though glucose is absolutely required. This is because glucose is utilized for biosynthetic rather than bioenergetic purposes, with pyruvate as the major nutrient utilized for energy production (Chi et al., 2020). As the embryo transitions from the morula to the blastocyst stage, glucose uptake increases sharply (Leese and Barton, 1984). This fact led us to investigate whether glucose begins to contribute to the TCA cycle in blastocysts and thereby becomes a primary bioenergetic resource. We traced the fate of ^13^C labeled glucose, comparing the 2C, morula and blastocyst stages. A combined use of UHPLC-MS and ion capillary chromatography-MS enables us to measure metabolite levels with a relatively small number of embryos (250 per sample compared with more than 1 million cells used in typical cancer studies). We complement these metabolic analyses with transcriptional profiling of the enzymes that are involved in these processes using RNA-seq and PCA analysis of either the entire transcriptome, or using only metabolic genes. In either case, we detect an inverted U-shaped trajectory in the gene expression profile as embryos progress from the early 2C stage to the blastocyst (Figure 1B, C). The smooth transition across the PCA projection axes indicates concerted changes in the gene expression profile of the embryo as development proceeds. The similarity in the overall profile for metabolic genes to the entire genome demonstrates that metabolic enzymes are tightly regulated during this period and that metabolism is not a static feature that facilitates development, but is dynamically controlled at the transcriptional level.

In labeling experiments, embryos are isolated as zygotes, and incubated in a medium in which all 6 glucose carbons are labeled uniformly with ^13^C (abbreviated U^13^C glucose). Metabolites are extracted at the 2C (2C; 48h), morula (M; 8-16 cells, 78h), and blastocyst (B; 32-64 cells, 96h) stages. Please note that unless stated otherwise throughout this paper the number of cells in 2C, morula and blastocyst stages are as described above with the hours referring to post-hCG (human chorionic gonadotropin) injection. The contribution of glucose can be distinguished from that of the other nutrients that are unlabeled (^12^C). The levels and the isotopologue distribution of labeled metabolites are then determined.

We find that the levels of glycolytic intermediates are highly dynamic during the transition from the 2C stage embryo to the blastocyst, and several of the glycolytic metabolites (for example, G6P, F6P, F16BP, G3P, and 3PG) increase significantly in abundance by the blastocyst stage (Figure 1D). In sharp contrast to this trend, a direct product of glycolysis, glycerol-3-phosphate (glycerol-3P), decreases by more than 10-fold during blastocyst formation (Figure 1D). Transcriptomic analysis shows that many glycolytic genes also peak in their expression at the blastocyst stage. Indeed, expression of genes such as *Hk2, Aldoa, Gapdh, Tpi1, Eno1*, hardly detectable in cleavage stage embryos, increase dramatically at the blastocyst stage, as does the expression of the glucose transporter *Slc25a3* (Figure 1E, Figure S1B, C). Of the glucose metabolism related genes, a smaller subset is much more highly expressed in early embryos than in the blastocyst. Prominent amongst them are isoforms of lactate dehydrogenase (*Ldhb* and *Ldhc*) and bisphosphoglycerate mutase (*Bpgm*) (Figure 1F). These genes encode enzymes that function in reactions that are linked to the bioenergetic status of the embryo, and do not function in the core glycolytic chain.

A significant advantage of using ^13^C labeling is that the breakdown of a nutrient can be traced by isotopologue analysis that detects the number of carbons that are derived from a labeled nutrient, in this case, labeled glucose. Thus, if a 6-carbon target metabolite is fully labeled by U^13^C glucose its mass will increase by 6 Da, and this peak is called the M6 isotopologue (Buescher et al., 2015). Whereas, if a 5-carbon metabolite is fully labeled, its mass will increase by 5 Da, yielding the M5 isotopologue. An unlabeled metabolite is designated M0 as it contains only ^12^C carbons. At the 2C and morula stages, M0 is the only isotopologue of TCA cycle metabolites that is detected when U^13^C glucose is used (Figure 1G). Close to 100% of citrate is M0 at the 2C and morula stages (Figure 1G). However, at the blastocyst stage, although 80% of the citrate molecules still remain M0, we start detecting low levels of M1-M3 peaks (18%) but M4-M6 isotopologues remain at less than 1% of the pool (Figure 1G). Partially labeled products are generally attributed to combinations of labeled and unlabeled metabolites generated during turnover of the TCA cycle. The rest of the detectable TCA cycle metabolites show a pattern of labeling that is similar to that seen for citrate (Figure 1G). Thus, glucose does not contribute to the TCA cycle metabolites at early stages of development, and it provides only minor amounts of carbons at the blastocyst stage. The contribution of glucose to acetyl-carnitine (derived from and reflecting the levels of acetyl-CoA) in the blastocyst continues to be low (10%) but at higher levels than at the morula stage (< 1%) (Figure S1A).

Another assay for evaluating glucose contribution to the TCA cycle is to determine its role in the formation of non-essential amino acids such as Ala, Gln, Glu and Asp. These are derived from metabolites such as pyruvate (for Ala), α-KG (for Glu/Gln) and oxaloacetate (for Asp).

Consistent with the results above, these non-essential amino acids are also not labeled by U^13^C glucose (Figure S1A). We conclude that there is a mechanism that prevents the majority of carbons from glucose entering the metabolite pools associated with the TCA cycle during all stages of preimplantation development.

Unlike for TCA cycle metabolites and amino acids, glucose plays an increasing role in populating nucleotides with its carbon. The contribution of glucose to nucleotides is barely detectable at the 2-cell stage. For example, unlabeled UMP (M0) is 97% in 2C embryos, 11% in morulae, and 6% in blastocysts. Similarly, unlabeled AMP (M0) is 93% in 2C embryos, 32% in morulae, and 9% in blastocysts. In the blastocyst, nearly all of the ribose carbons in nucleotides are derived from exogenously provided glucose (Figure 1H).

The prominent glucose derived nucleotide isotopologue is M5, accounting for the 5 labeled carbons of the ribose sugar, but not higher indicating that the nucleobases are not labeled by glucose. Consistent with this notion, the glycine pool, required for *de novo* purine nucleobase synthesis, does not derive labeled carbons from glucose (Figure S1A). This allows us to conclude that rather than exogenously provided glucose, an endogenous source, such as maternal glycine or salvage of bases, is required for nucleobase formation. This is confirmed by labeling experiments in which U^13^C glycine is provided to the embryos in the presence or absence of (unlabeled) glucose. In the presence of glucose, U^13^C glycine contributes to the generation of M2-M4 isotopologues of purine nucleotides. However, in media in which glucose is absent, no labeled purine nucleotide isotopologue is detected, demonstrating that ribose from exogenous glucose is required for *de novo* purine synthesis (Figure 1I). Thus, purine nucleobases are derived in part from endogenous glycine, whereas ribose sugars are entirely derived from exogenously provided glucose.

A majority of the purine and pyrimidine biosynthetic genes increase in expression as development proceeds. Genes such as *Gart, Paics, Impdh, Pfas*, and *Atic* participate in multiple steps in purine biosynthesis (Lane and Fan, 2015) and they peak in expression at the morula and early blastocyst stages (Figure 1J, K). Pyrimidine synthesis genes, such as *Cad* and *Umps* that function in important steps in the *de novo* pathway, peak at the 4-8 cell stages (Figure 1L). Genes involved in the purine nucleotide cycle, *Adss, Adsl* and *Ampd3* show an opposite trend in that their expression peaks during the early stages and either declines (for *Adss* and *Ampd3*) or plateaus (for *Adsl*) later in development (Figure 1K, S1D). These genes help maintain a balanced adenine nucleotide pool and as byproducts of this cycle, they also generate fumarate and ammonia (Aragon et al., 1981).

Genes for purine degradation enzymes (*Xdh, Ada, Gda, Pnp, Pnp2*, and *Adk*) are more highly expressed early and decrease in expression as development proceeds (Figure 1M). Whereas genes encoding purine salvage enzymes, such as *Hprt* and *Aprt*, show an opposite trend, peaking at the late morula and early blastocyst stages (Figure S1E). Interestingly, hypoxanthine, which is formed by degradation of adenine, and is used by *Hprt* in the salvage pathway of nucleotide synthesis, is barely detectable in 2C embryos, and increases by nearly 100-fold in blastocysts (Figure S1F). These data imply that the capacity of the salvage pathway of purine nucleotide synthesis increases during the course of preimplantation development. In summary, nucleotide synthesis is predominantly a late preimplantation phenomenon and therefore the biosynthetic genes appear later in development. The genes encoding degradation enzymes on the other hand are more abundant at the earlier stages.

### Pyruvate and lactate metabolism during preimplantation development

We then evaluated the possible contribution of pyruvate/lactate to the TCA cycle. When U^13^C pyruvate and U^13^C lactate are provided in the medium, more than 90% (at 2C) and 83% (for blastocysts) of acetyl-carnitine (a surrogate for acetyl-CoA) is the M2 isotopologue (Figure 2A). M6 labeled citrate is 61% at the 2C stage with M0 at 6% of the pool. Intermediate isotopologues account for the remaining citrate (Figure 2A). Aconitate and α-KG show broadly similar labeling patterns as citrate (Figure 2A).

**Figure 2.**
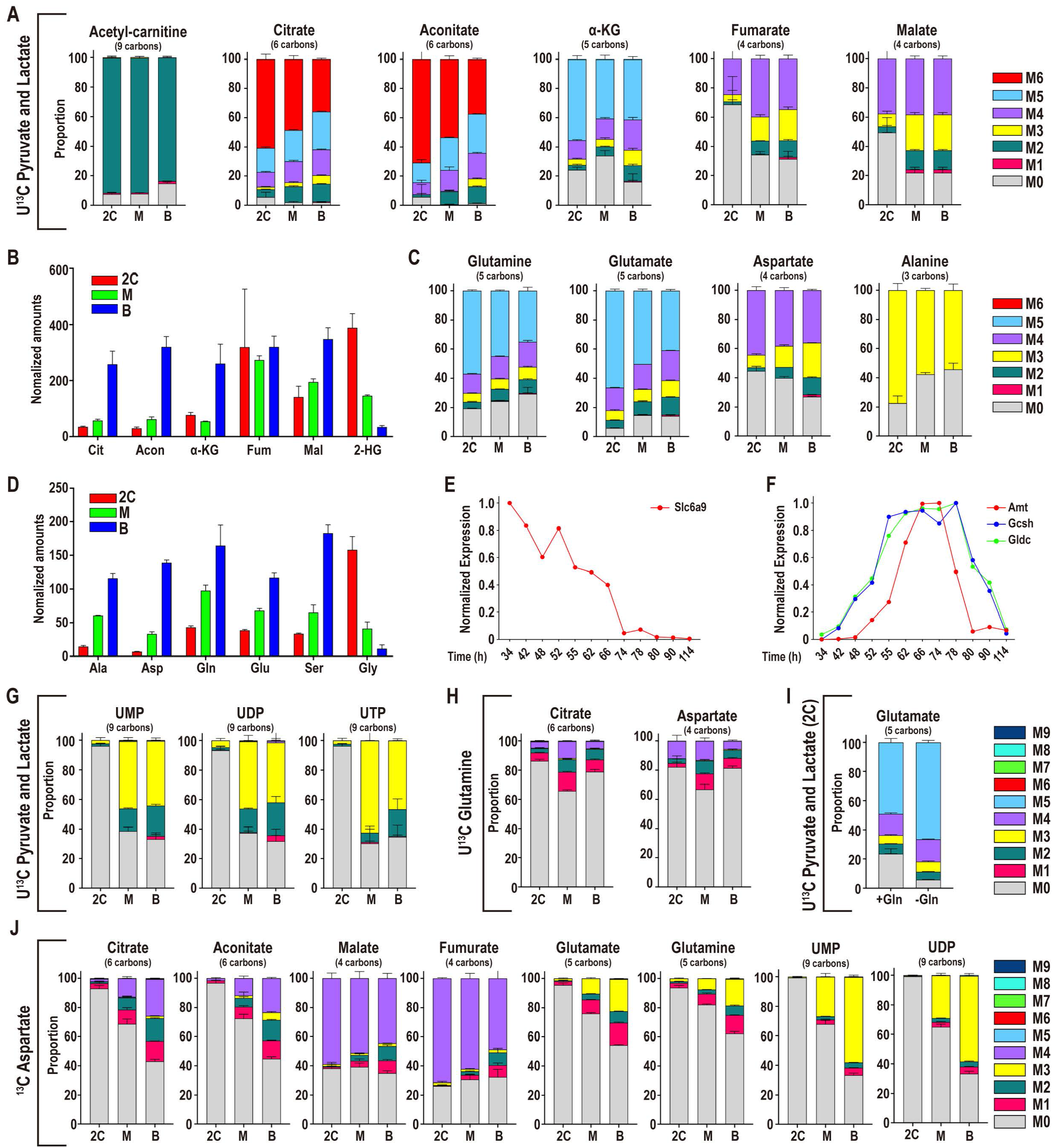
Pyruvate and lactate metabolism during preimplantation development. **(A-D)** U-^13^C pyruvate and U-^13^C lactate contributions to TCA cycle metabolites and amino acids. In all figure panels, stages are represented as 2C: 2-cell, M: morula, B: blastocyst. **(A)** Isotopologue contributions to acetyl-carnitine and the TCA cycle metabolites. For acetylcarnitine, M2 represents the acetyl group. Citrate and aconitate (Class I metabolites), but not fumarate and malate (Class II), are extensively labeled throughout the preimplantation period. **(B)** Scaled amounts of TCA intermediates at each stage. **(C)** Isotopologue contributions to the amino acids Glu, Gln, Ala and Asp. **(D)** Scaled amounts of non-essential amino acids at each stage. **(E, F)** The glycine transporter *Slc6a9* **(E)** peaks in gene expression earlier than the enzymes of the glycine cleavage system **(F)**. **(G)** U-^13^C pyruvate and U-^13^C lactate contribute to pyrimidine formation at later stages. **(H)** U-^13^C Gln only contributes minor amounts of carbon to the TCA cycle. **(I)** U-^13^C pyruvate and U-^13^C lactate generate a majority of Glu in 2C embryos both when Gln is present (+Gln) and absent (-Gln). **(J)** In 2C embryos, U-^13^C Asp contributes to Class II metabolites, but not to Class I metabolites. In blastocysts, the Asp contribution to Class I metabolites increases.

The proportion of fully labeled citrate decreases from 61% in 2C embryos to 36% in blastocysts. The decrease in M6 citrate is not caused by an increase in M0, which remains almost undetectable, but is associated with a rise in intermediate isotopologues. We conclude that at all stages pyruvate and lactate are the major players that populate acetyl-CoA, citrate, aconitate and α-KG (Figure 2A).

The labeling pattern of malate and fumarate, however, is markedly different from citrate and aconitate (Figure 2A). For example, at the 2C stage, only 37% of malate is fully labeled (M4), and 50% is M0. The amount of malate that is M0 decreases to 22% in blastocysts compared to 50% in 2C embryos. Intermediate isotopologues, such as M3, rise, and account for the changes (M3 is 9% in 2C, 25% in blastocysts). Fumarate shows similar labeling patterns as malate (Figure 2A).

We find that between the 2C and blastocyst stages, while citrate, aconitate and α-KG, all increase by about 10-fold, malate and fumarate levels increase only by about 2.5-fold (Figure 2B). These data are consistent with our published model that citrate, aconitate and α-KG (that we call Class I metabolites) behave differently from malate and fumarate (Class II metabolites) in their sensitivity to and recovery from nutrient starvation (Nagaraj et al., 2017). In conclusion, even at the blastocyst stage, Class I and Class II metabolites produced from the two different halves of the TCA cycle do not mix freely in the embryo. Class I metabolites are primarily derived from pyruvate and lactate, while the Class II pools have a large unlabeled component, which must be generated from an endogenous source. The disequilibrium between the metabolites derived from the right and left halves of the TCA cycle is most marked at the 2C stage and relaxes somewhat in the blastocyst.

We next determined the contribution of pyruvate and lactate to the formation of amino acids. At the 2C stage, only four amino acids (Ala, Asp, Gln, Glu) are labeled by pyruvate and lactate (Figure 2C). Amongst these, Glu and Asp are derived from direct products of the two different arms of the TCA cycle. Glu (derived from α-KG) is only 5% unlabeled, while as much as 45% of Asp (derived from malate) is unlabeled (Figure 2C). This is consistent with the concept of a disequilibrium between Class I and Class II metabolites of the TCA cycle.

Other amino acids such as Pro, Arg and Asn that could, in principle, be generated from pyruvate and lactate, are not derived from these nutrients and must necessarily be derived from endogenous sources (Figure S2A). Interestingly, several amino acids increase dramatically in abundance as development proceeds, although none is provided in the medium. For example, Tyr, Phe, and Trp, all increase by more than 10-fold, and Asp by 20-fold, between the 2C and blastocyst stages (Figure 2D, S2B). The simplest explanation is that essential amino acids are derived from increased degradation of endogenous proteins as development proceeds.

Gly and Thr do not increase during development (Figure 2D, S2B), likely attributable to their rapid consumption as the embryo matures. Thr levels are highest at the morula stage, coincident with a peak in the expression of threonine dehydrogenase (*Tdh*) (Figure S2C). Glycine is highest in 2C embryos and decreases by 14-fold in blastocysts (Figure 2D, see also (Baltz, 2001)). The early stage elevated glycine levels can in part be accounted for by the high expression of the glycine transporter, *Slc6a9*, in oocytes and zygotes prior to our embryo isolation (Figure 2E, (Steeves et al., 2003)). Following ZGA, the expression of the three enzymes of the glycine cleavage system (GCS) increases (Figure 2F), which correlates with a fall in glycine levels.

We next determined the role of pyruvate and lactate in nucleotide formation using ^13^C labeling. For both purines and pyrimidines, the ribose moiety is generated from glucose (Figure 1H). Isotopologue analysis shows that pyruvate and lactate do not contribute carbon to purine nucleobases at any stage (Figure S2D). For pyrimidines, this holds true at the 2C stage. However later, at the morula stage, there is increased labeling, suggestive of new nucleotide synthesis (Figure 2G). For example, UMP is 38% M0, and 61% M2/M3. Other pyrimidine nucleotides show a similar labeling pattern to UMP, and, in general, at later stages of development, pyrimidine nucleotides have a significant contribution from pyruvate and lactate (Figure 2G). In summary, until ZGA, the embryos rely on maternally deposited nucleotides, and with expression of new enzymes following the 2C stage, *de novo* nucleotide synthesis gains prominence. At this point exogenously provided glucose generates the ribose moiety of nucleotides, while pyruvate-derived and endogenous metabolites combine to generate nucleobases, and together they form newly synthesized nucleotides.

In cancer cells, glucose often plays a greater role in glycine and nucleobase synthesis than we find in the embryo, and also, exogenous Gln is taken up in large amounts and its carbons are used to synthesize Asp (Birsoy et al., 2015; Sullivan et al., 2015). In the embryo, Gln and Asp, that are critical for multiple biosynthetic reactions, are both formed from pyruvate (Figure 2C), although its contribution to Gln is greater than it is to Asp (Figure 2C). Following uptake into the cell, Gln is converted to glutamate by the enzyme glutaminase. Both glutaminase isoforms (*Gls* and *Gls2*) are expressed in the embryo, peaking in expression at the 4-8C stage (Figure S2E). To determine a possible contribution of Gln to central carbon metabolites during development, we cultured embryos in normal medium supplemented with 1mM U^13^C Gln and find that the predominant TCA cycle intermediate isotopologue at each of the developmental stages examined is M0 (i.e., not labeled by Gln) (Figure 2H, S2F). Gln is also not a significant carbon source for Asp or pyrimidine nucleotides in the preimplantation embryo (Figure 2H, Figure S2F). These results are qualitatively different from the high rates of glutamine oxidation and incorporation seen in cancer cells (Hensley et al., 2013). Moreover, and perhaps surprisingly, glutamate, which in other systems is largely derived from glutamine, continues to be resourced by pyruvate even when glutamine is included in the media (Figure 2I). Furthermore, added Gln cannot rescue the block in development seen upon inhibition of GLUL (glutamine synthetase) that converts Glu to Gln and is expressed at peak levels at the 4-8 cell stage (Figure S2E, G). Thus, since Glu is formed from pyruvate (via α-KG), Gln in the embryo is indirectly derived from pyruvate by conversion of Glu to Gln using the essential enzyme GLUL.

Like Gln, the amino acid Asp also has a number of critical biosynthetic roles within a cell. In labeling experiments in which U^13^C Asp is added to the medium, we find that pyrimidine nucleotides, TCA cycle metabolites, and Glu and Gln receive Asp carbons in a stage specific manner (Figure 2J). At the 2C stage, added U^13^C Asp hardly labels pyrimidine nucleotides whereas in later stage embryos, it readily does so, such that in blastocysts, about 60% of UMP is labeled (Figure 2J, S2H). Furthermore, in the 2C embryo, U^13^C Asp efficiently labels malate (unlabeled M0 = 38%) and fumarate (M0 = 26%) but not citrate (unlabeled M0 = 93%), aconitate (M0 = 97%), or Glu (M0 = 95%) and Gln (M0 = 94%) (Figure 2J). In blastocysts, Asp can contribute significantly more to citrate (M0 = 43%), aconitate (M0 = 45%), Glu (M0 = 54%) and Gln (M0 = 62%) (Figure 2J). These results further substantiate the pyruvate labeling data (Figure 2A) and emphasize that at the 2C stage there exists a disequilibrium between the Asp-malate-fumarate and the citrate-aconitate-αKG arms of the TCA cycle. This disequilibrium is attenuated as development progresses to the blastocyst stage. This change is one more indication of increased metabolic plasticity with developmental progression.

### Adaptation to nutrient conditions

Thus far, all experiments described involve normal culture conditions, but historically, individual nutrient deprivation experiments have informed our current understanding of metabolic requirements (Brown and Whittingham, 1991). The classic example of diversification of nutrient requirement is that embryos must receive pyruvate before, but not necessarily after, the 2C stage in order to progress further (Biggers et al., 1967; Lane and Gardner, 2005). Our PCA analysis shows that pyruvate omission from the zygote stage, causes a clear retardation in the expression of developmentally regulated metabolic genes, while deprivation from the late (50h) 2C stage does not substantially affect their trajectory (Figure 3A, A’ Figure S3A, A’). It is clear that a study of the expression of pathway-specific enzymes and metabolites rather than of global changes in the entire metabolic repertoire is necessary to meaningfully correlate metabolomic and transcriptomic profiles with phenotypic analysis.

**Figure 3.**
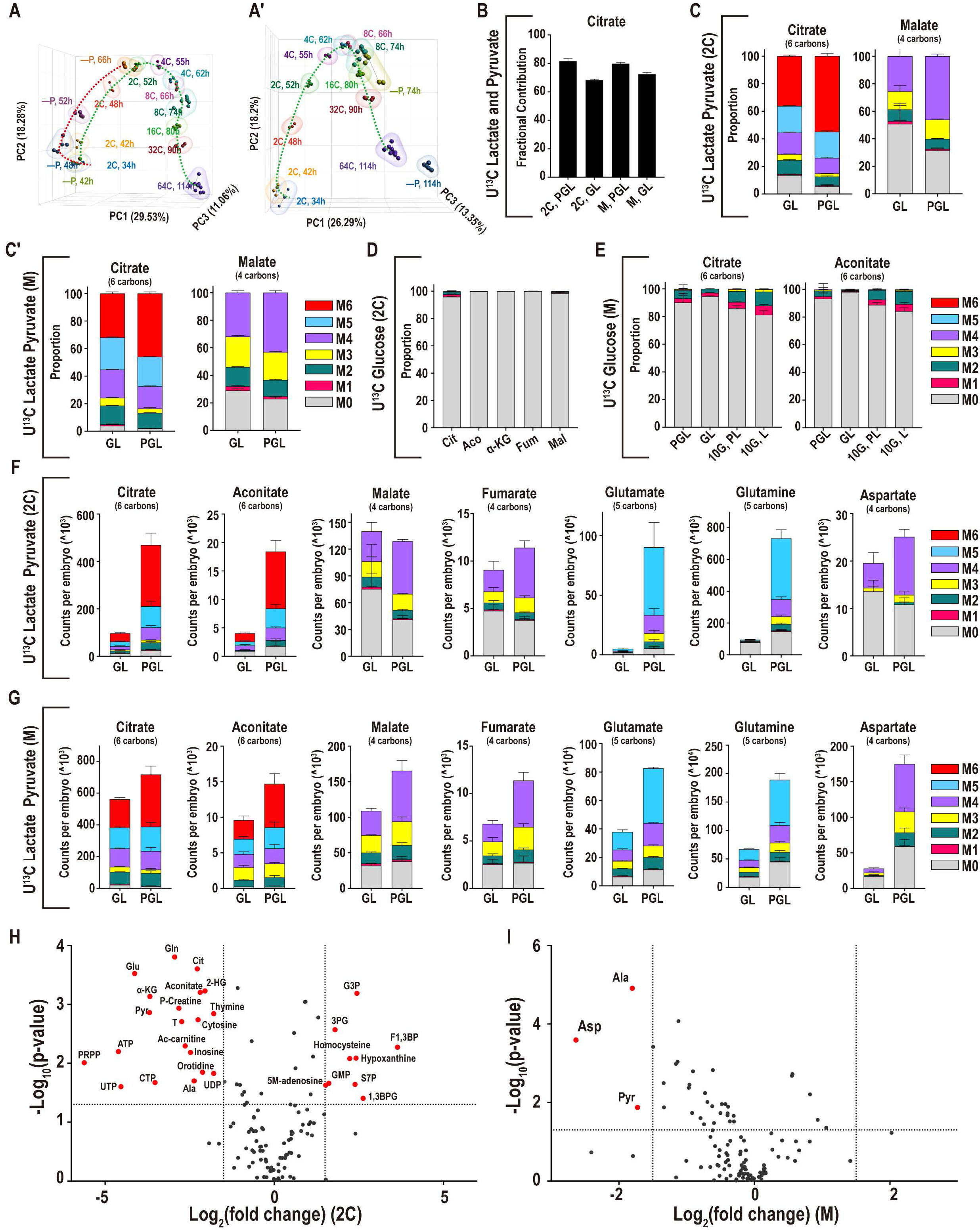
Adaptation to nutrient conditions. **(A-A’)** PCA analysis of genes that are directly involved in metabolic processes (1947 genes) in embryos developing with and without pyruvate. Pyruvate is removed from the zygote stage **(A)** or the late 2C stage **(A’)**. **(B)** In both 2C embryos and morulae, U-^13^C pyruvate/lactate contributes a majority of carbons to the total citrate pool in the presence (PGL) or absence (GL) of pyruvate. **(C, C’)** At both the 2C **(C)** and morula **(C’)** stages, the isotopologues of citrate that are formed from U-^13^C lactate alone (no pyruvate is present, GL) are similar to the isotopologues that are formed when both U-^13^C pyruvate and U-^13^C lactate (PGL) are present. **(D, E)** The contribution of U-^13^C glucose to the TCA cycle metabolites remains very low in 2C embryos **(D)** as well as in morulae **(E)** in GL media (lacking pyruvate), even when a high amount of glucose (10mM instead of the normal 0.2mM) is provided (10G, L). **(F, G)** Formation of Class I metabolites from U-^13^C lactate/pyruvate decreases when pyruvate is withheld from 2C embryos **(F)**, but less so in morulae **(G)**. Fumarate and malate are less sensitive, and Asp only decreases significantly in morulae. **(H)** In 2C embryos 20 metabolites decrease significantly, and 9 metabolites increase upon pyruvate withdrawal (GL). **(I)** In morulae, only Asp, pyruvate, and Ala decrease significantly. Metabolites are highlighted if P < 0.05 and the log_2_ fold-change > +/-1.5.

In ^13^C labeling experiments to determine nutrient utilization during pyruvate withdrawal, embryos are cultured in either 1) U^13^C lactate, unlabeled glucose and no pyruvate, or 2) U^13^C glucose, unlabeled lactate and no pyruvate. Comparisons are made to corresponding controls in which both U^13^C pyruvate and U^13^C lactate are provided. At both the 2C and morula stages, lactate continues to contribute a majority of carbons to the TCA cycle when pyruvate is withheld (unlabeled citrate: M0 = 14% in 2C, M0 = 4% in morula). At both the 2C and morula stages, when pyruvate is not present, the contribution of glucose to the TCA cycle remains very low (unlabeled citrate: M0 = 95% in 2C, M0 = 96% in morula), even if glucose is increased by 50-fold over normal medium levels (Figure 3B-E, Figure S3B-E). It is clear that pyruvate withdrawal does not cause a switch to glucose-based bioenergetics, instead the TCA cycle continues to be driven by pyruvate that is intracellularly derived from imported lactate (Figure S3F, G).

The pyruvate withdrawal experiments also highlight the disequilibrium between the two halves of the TCA cycle and the increase in metabolic plasticity during development, features of embryo metabolism that were discussed earlier in the context of normal development. The distinction is most clear for 2C embryos. For example, upon pyruvate withdrawal, total citrate decreases by 4.8-fold (in 2C) but only 1.3-fold (in morulae) and the fully labeled M6 citrate falls by 7.4-fold (in 2C) but also only 1.8-fold (in morulae) (Figure 3F, G). Similarly, aconitate decreases by 4.6-fold (in 2C) but 1.5-fold (in morulae), and M6 aconitate decreases by 6.9-fold (in 2C) and 2.3-fold (in morulae). In contrast, Class II metabolites behave quite differently; malate and fumarate actually increase by 1.1-fold, and 1.3-fold respectively in 2C embryos upon pyruvate withdrawal, whereas, in morulae, malate and fumarate decrease by 1.5-fold and 1.7-fold (Figure 3F, G). Metabolites that are associated with energy status, such as phosphocreatine, ATP, and acetyl-carnitine, also decrease in 2C embryos, but are unaltered in morulae, and a number of metabolites involved in pyrimidine nucleotide metabolism decrease in 2C embryos but are not sensitive to pyruvate withdrawal in later stage embryos (Figure 3H, I).

The most striking decrease in 2C embryos is for the amino acids Gln (8-fold decrease) and Glu (17-fold decrease), whereas for morulae this decrease is less dramatic, Gln (2.8-fold) and Glu (2-fold) (Figure 3F, G). At the 2C stage pyruvate withdrawal causes a huge drop in fully U^13^C lactate labeled Glu (M5, 32-fold) and Gln (M5, 67-fold), while the corresponding drops in morulae are minor, Glu (M5, 3-fold) and Gln (M5, 4-fold). Thus, while the synthesis of Glu and Gln is sensitive to pyruvate levels at both stages, 2C embryos are far more sensitive than morulae. Altogether, the above results show that 2C embryos suffer severely from nutrient depletion when pyruvate is withdrawn, whereas morulae are far more resistant. This result is most obvious when expressed as a volcano plot of fold-differences and their P-values (Figure 3H, I).

### Mechanisms of metabolic plasticity

Asp is the only metabolite that decreases in morulae (6-fold), while being stably maintained in 2C embryos upon pyruvate withdrawal (only a 1.2-fold decrease) (Figure 3F, G). The dramatic fall in Asp levels in morulae, but not 2C embryos, in response to pyruvate withdrawal is particularly interesting, because Asp levels also increase dramatically as development proceeds under normal growth conditions, with the levels of Asp 20-fold greater in blastocysts than in 2C embryos (Figure 2D). A number of studies have shown that Asp is highly sensitive to the redox state of the cell (Birsoy et al., 2015; Sullivan et al., 2015), and Asp has an important role in the malate-aspartate (Mal-Asp) shuttle, which controls NADH/NAD^+^ levels in the cytoplasm. Thus, the large variance in Asp levels during normal development may signal changes in NADH:NAD^+^ control mechanisms. In the Mal-Asp shuttle, MDH interconverts malate and oxaloacetate, and the latter then interconverts with Asp by the action of the aspartate aminotransferase enzyme GOT (Lane and Gardner, 2005) (Figure 4F). The increase in Asp, during development or when pyruvate is withheld, indicates a decrease in NADH, because when oxaloacetate is converted to malate, NADH is consumed and NAD^+^ is produced.

**Figure 4.**
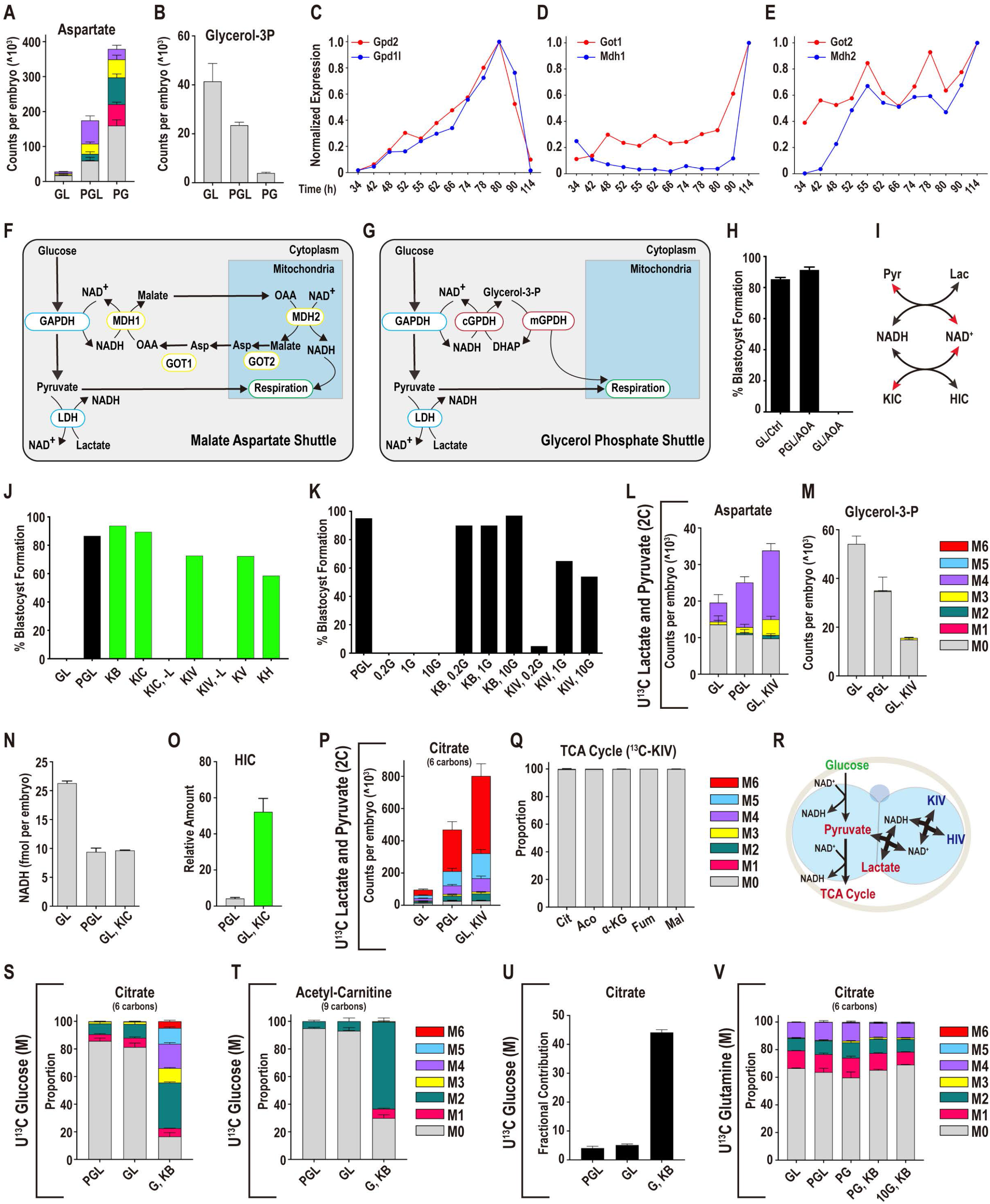
Mechanisms of metabolic plasticity. **(A)** Asp increases when lactate is absent (PG), and decreases when pyruvate is withheld (GL). **(B)** In contrast, glycerol-3-phosphate (glycerol-3P) decreases when lactate is absent and increases when pyruvate is withheld. **(C)** The glycerol-3P shuttle components *Gpd1l* and *Gpd2* that are expressed in the embryo peak at the morula stage. **(D)** The cytoplasmic Mal-Asp shuttle components, *Mdh1* and *Got1*, peak in expression in blastocysts, and **(E)** the mitochondrial components *Mdh2* and *Got2* increase in expression from the late 2C stage. **(F, G)** Schematic of the Mal-Asp shuttle **(F)** and the glycerol-3P shuttle **(G)**. **(H)** Following the 2C stage, inhibition of the Mal-Asp shuttle using 0.5 mM AOA does not impact the development of embryos cultured with pyruvate, lactate and glucose (PGL), but blocks development of embryos cultured in GL media **(I)** Schematic showing how pyruvate and lactate, and α-KIC and HIC, control NADH and NAD^+^ levels. **(J)** Alternative α-ketoacids (1mM), allow zygotes to develop into blastocysts in the absence of pyruvate, as long as lactate is provided. In a medium lacking pyruvate (GL), embryos do not form blastocysts unlike embryos that are cultured with pyruvate (PGL). The conditions represented by green bars all lack pyruvate (-P). The α-ketoacids KB, KIC, KIV, KV and KH all rescue (-P) embryos unless both pyruvate and lactate are absent (KIC, -L) (KIV, -L). Glucose (G) is present in all samples above, and pyruvate is absent in all samples except control (PGL). **(K)** Embryos that are transferred into pyruvate and lactate free media at the late 2C stage (after ZGA) (50h) are not viable over a range of glucose concentrations (0.2mM (0.2G), 1mM (1G) or 10mM (10G)) unless an α-ketoacid, such as KB or KIV, is provided. **(L, M)** In 2C embryos, aspartate decreases and glycerol-3P increases in GL (U-^13^C lactate, U- ^12^C glucose) media. KIV fully restores Asp levels and decreases the amount of glycerol-3P. The PGL (U-^13^C pyruvate and U-^13^C lactate) and GL (U-^13^C lactate) controls are shared with Figure 3F as these experiments were performed at the same time. **(N)** NADH levels in zygotes with pyruvate, lactate and glucose (PGL), glucose and lactate (GL), or GL supplemented with 1mM KIC (GL, KIC). **(O)** Hydroxycaproate (HIC) increases in embryos that are provided with KIC. **(P)** U-^12^C KIV restores citrate levels in 2C embryos deprived of pyruvate by facilitating the use of U-^13^C lactate. The PGL (U-^13^C pyruvate and U-^13^C lactate) and GL (U-^13^C lactate) control samples are shared with Figure 3F, because these experiments were performed at the same time. **(Q)** U-^13^C KIV does not contribute carbon to the TCA cycle in 2C embryos cultured in GL media and therefore does not act as a nutrient but functions as a redox balancer. **(R)** Schematic illustrating how α-KIV facilitates the conversion of lactate to pyruvate. α-KIV raises NAD^+^ levels via LDHC. This NAD^+^ allows LDHB to convert lactate into pyruvate. **(S, T)** 10mM U-^13^C glucose contributes insignificant amounts of carbon to citrate **(S)** and acetylcarnitine **(T)** in morulae in either PGL or GL media. In contrast, glucose contributes substantially with both pyruvate and lactate omitted if α-KB is provided. The control samples are shared with Figure 3E, because these experiments were performed at the same time. **(U)** For embryos cultured in PGL or GL, 10mM U-^13^C glucose contributes a minor amount of carbon to the total citrate pool. But with both pyruvate and lactate omitted, and α-KB provided, glucose contributes substantially to citrate. **(V)** U-^13^C glutamine contributes minor amounts of carbon to citrate even when both pyruvate and lactate are omitted from the medium and unlabeled 10mM glucose and α-KB are provided.

A related shuttle, called the glycerol-3P shuttle, is able to fulfil a similar role to the Mal-Asp shuttle (Figure 4G). We find that under normal culture conditions, glycerol-3P, which functions in the glycerol-3P shuttle, falls 10-fold between the 2C and blastocyst stages (Figure 1D). The decrease in glycerol-3P also signals a decrease in NADH, since conversion of dihydroxyacetone phosphate (DHAP) to glycerol-3P consumes NADH and generates NAD^+^. Pyruvate deprivation will cause a further rise in the NADH:NAD^+^ (Veech et al., 2019). This is borne out by the fact that Asp levels are 13-fold lower and glycerol-3P is 10-fold higher in morulae when they are cultured in a pyruvate-free medium (with normal lactate and glucose) compared with one in which lactate is depleted (with normal pyruvate and glucose) (Figure 4A, B).

Gene expression analysis supports the metabolomic data in that the enzymes that function in these two shuttles display a dynamic expression profile. For example, *Gpd1l* and *Gpd2* (glycerol-3P shuttle) increase by more than 50-fold between the 2C and morula stages (Figure 4C). Similarly, for the Mal-Asp shuttle, *Mdh1* and *Mdh2* increase by approximately 20-fold and 4-fold respectively, and *Got1* and *Got2* enzymes increase by 9-fold and 2.5-fold respectively, between the 2C and blastocyst stages (Figure 4D, E, S4A).

In a medium replete with pyruvate, GOT (and therefore the Mal-Asp shuttle) inhibition by AOA (aminooxyacetic acid) has no phenotypic consequence for the growth of the embryo (Figure 4H). However, pyruvate deprived embryos treated with this inhibitor block in development at the morula stage (Figure 4H, see also (Lane and Gardner, 2005)). In summary, while the protein and metabolic components of the Mal-Asp shuttle increase in later stages of development, an essential role for this shuttle to generate cytoplasmic NAD^+^ is phenotypically apparent only upon pyruvate withdrawal.

In contrast to the elevated expression of the shuttle components at later stages, at the 2C stage, by far the most important dehydrogenase that is expected to control the redox balance of the embryo is LDHB. Conversion of lactate to pyruvate by LDHB leads to high NADH levels (Veech et al., 2019). LDHB activity in the early embryo is 10-fold greater than in any somatic tissue, and its measured activity far exceeds that of all other metabolic enzymes (Brinster, 1965; Epstein et al., 1969). As development proceeds, we detect a striking 2000-fold decrease in the LDHB mRNA level (Figure 1F). We surmise that the changes in relative levels of the dehydrogenases that control NADH:NAD^+^ balance is an important determinant of how NADH:NAD^+^ impacts, and is impacted by, broader cell metabolism during development. This will affect the flow of carbons between major metabolic pathways. It is reasonable to propose that this change in redox control during normal development is at the core of the increase in metabolic plasticity that arises as the embryo advances through the preimplantation process.

### Rebalancing redox ratios during early stages of development

The above premise is testable by using various α-ketoacids, some naturally occurring and others not, to mimic the effects of pyruvate as a substrate of LDH. Pyruvate (α-ketopropionate) is the simplest of all α-ketoacids that have a direct linkage between a ketone and a carboxyl functional group. A number of studies have shown that a mimic such as α-KB (α-ketobutyrate) is readily taken up by cells and reduced by LDHB (Sullivan et al., 2015). This process generates NAD^+^ and α-hydroxybutyrate, and provides a way to alter the redox status of the cell. Another isoform of LDH, LDHC, also present in the embryo (Figure 1F, (Coonrod et al., 2006)) is highly promiscuous and is able to use α-KB, but also a wider range of α-ketoacids, such as α-KIV (α-ketoisovalerate) and α-KIC (α-ketoisocaproate), which are not substrates for LDHB. Even α-ketoacids, such as α-KV (α-ketovaleric acid), that have no known function in mammalian cells, are effective substrates for LDHC. In biochemical assays such α-ketoacids are reduced by LDHC, and they generate NAD^+^ and their corresponding α-hydroxyacids (Blanco et al., 1976). Thus, in principle the presence of LDHC affords us the capability to modulate NAD^+^ and NADH independently of the lactate/pyruvate interconversion that is dominated by LDHB.

When pyruvate deprived zygotes (1C) are provided with a range of α-ketoacids, α-KB, α-KIV, α-KIC, α-KV, or α-KH, they no longer block at the 2C stage and are able to make blastocysts (Figure 4J). α-ketoacids cannot rescue development if both pyruvate and lactate are absent (Figure 4J, marked KIC, -L and KIV, -L). The rescue by multiple α-ketoacids that are known substrates of LDHC lends credibility to the idea that rebalancing redox is an important mechanism underlying developmental progression.

Unlike at the earlier stage, embryos develop to blastocysts even when lactate and pyruvate are both withdrawn from the medium as long as both glucose and an α-ketoacid are provided. Embryos that are provided with only glucose but no α-ketoacid die rapidly. Late (post-ZGA) 2C embryos provided with glucose and α-KB develop into blastocysts at a rate that is comparable to that of control embryos (Figure 4K). α-KIV is also able to restore development, but requires higher amounts of glucose.

Multiple lines of evidence establish that α-ketoacids provided to preimplantation embryos allow them to generate NAD^+^ and consume NADH. First, when embryos are provided with, for instance, α-KIV, the changes in Asp and glycerol-3P levels that are caused by pyruvate withdrawal are fully rescued (Figure 4L, M). In 2C embryos, the amount of fully labeled (M4) Asp is nearly 4-fold greater in α-KIV treated embryos, compared with embryos that are only provided with U^13^C lactate (Figure 4L). Second, providing α-ketoacids to the embryo prevents the increase in total NADH that is observed when pyruvate is withheld (Figure 4N), as assessed by cycling assays. For example, the more than 2-fold increase in NADH levels caused by pyruvate withdrawal in 2C embryos is reverted upon addition of α-KIC to the normal level that is seen in the presence of pyruvate (Figure 4N). Finally, when an α-ketoacid is included in the medium a high level of the corresponding α-hydroxyacid is detected. For example, in the presence of α-KIC the level of its α-hydroxyacid, α-HIC, which can only form in a redox related reaction, increases by more than 12-fold (Figure 4O).

The above findings are in complete agreement with labeling data that show that when pyruvate-deprived embryos are provided with U^13^C lactate and U^12^C α-KIV, lactate continues to contribute a majority of carbon to the TCA cycle and its associated amino acids (Figure 4P, Figure S4B-D). Indeed, Figure 4P shows that under conditions of pyruvate withdrawal, α-KIV treated embryos have 14-fold more M6 citrate labeled by lactate compared to embryos not provided with an α-ketoacid, and the rescue with α-KIV results in even more citrate than that seen with pyruvate. Importantly, ^13^C α-KIV does not label the TCA cycle (Figure 4Q). Thus, α-KIV acts by facilitating lactate to pyruvate conversion, and is not by itself a nutrient source for the embryo. As a whole, these results support the idea that α-ketoacids rescue development by a redox mediated mechanism. We conclude that at all development stages, lactate can substitute for pyruvate, but only if the cytoplasmic redox state is correctly balanced (Figure 4R).

Normally U^13^C glucose does not label TCA cycle components and acetyl-CoA (Figure 1G). If pyruvate is present, even with 10mM labeled glucose provided in the medium, more than 86% of citrate is M0 (unlabeled). Similarly, if pyruvate is absent, greater than 81% of citrate is unlabeled (Figure 4S). Expressed in a different way, glucose contributes only 4% (with pyruvate present) or 5% (with pyruvate absent) of the total number of citrate carbons (Figure 4U). Similarly, under these conditions the acetyl group of acetyl-CoA (reported by the M2 peak of acetyl-carnitine) is labeled 5% (with pyruvate present) and 7% (without pyruvate) (Figure 4T). Strikingly, with pyruvate and lactate both absent but with α-KB provided for redox balance, U^13^C glucose becomes a significant donor of carbons to the TCA cycle and to acetyl-CoA (Figure 4S, T, S4E). 83% of the citrate pool is now either fully or partially labeled by carbons from glucose. Similarly, 63% of the acetyl-CoA (M2 acetyl-carnitine) is labeled by glucose (Figure 4T). Thus, glucose does not contribute to the TCA cycle when pyruvate and lactate are present, and embryos are non-viable when provided with glucose alone. Embryos can survive with lactate and glucose alone, but then lactate provides carbon to the TCA cycle. For glucose to be an effective nutrient to the TCA cycle, not only do pyruvate and lactate need to be absent, but the cell needs to rebalance its redox utilizing the provided α-ketoacid.

Even with a redox balancer added, glucose is not as efficient as pyruvate/lactate in providing carbons for bioenergetic purposes (Figures 2A, 4T, U). With redox rebalance, glucose derived carbons mix with carbons from unlabeled sources that enter the TCA cycle pool as acetyl-CoA groups and perhaps at other anaplerotic entry points. This is an example of the expansion of the metabolic repertoire through development and/or during stress. In other systems, such as ESCs and cancers, Gln plays this role, but that is not true for the embryo since the contribution of Gln to the TCA cycle remains unchanged under these conditions (Figure 4V, S4F, G).

### Further metabolic plasticity in later stage embryos

Late 2C (50h) or early 4C (54h) stage embryos are non-viable when both pyruvate and lactate are removed from the medium and embryos do not survive using glucose alone (Figure 5A). However, we find that when transferred to a glucose only medium at 60h (mid-4C), 80% of the embryos are able to develop into blastocysts, and when transferred at the uncompacted 8C stage (68h), all the embryos that are grown without pyruvate and lactate develop into blastocysts (Figure 1A, 5B). Therefore, as development proceeds, the nutrient requirement becomes more flexible such that by the 8C stage, glucose alone is sufficient to sustain development even in the absence of an α-ketoacid. Remarkably, at the 8-16C stage (76h), 70% of the embryos are able to develop into blastocysts without the provision of any nutrients from the environment (Figure 5C). Following 86 hours (late morula, ~32C), 100% of the embryos form fully expanded blastocysts (114h) without any exogenously provided nutrients: pyruvate, lactate and glucose (Figure 1A, 5C).

**Figure 5.**
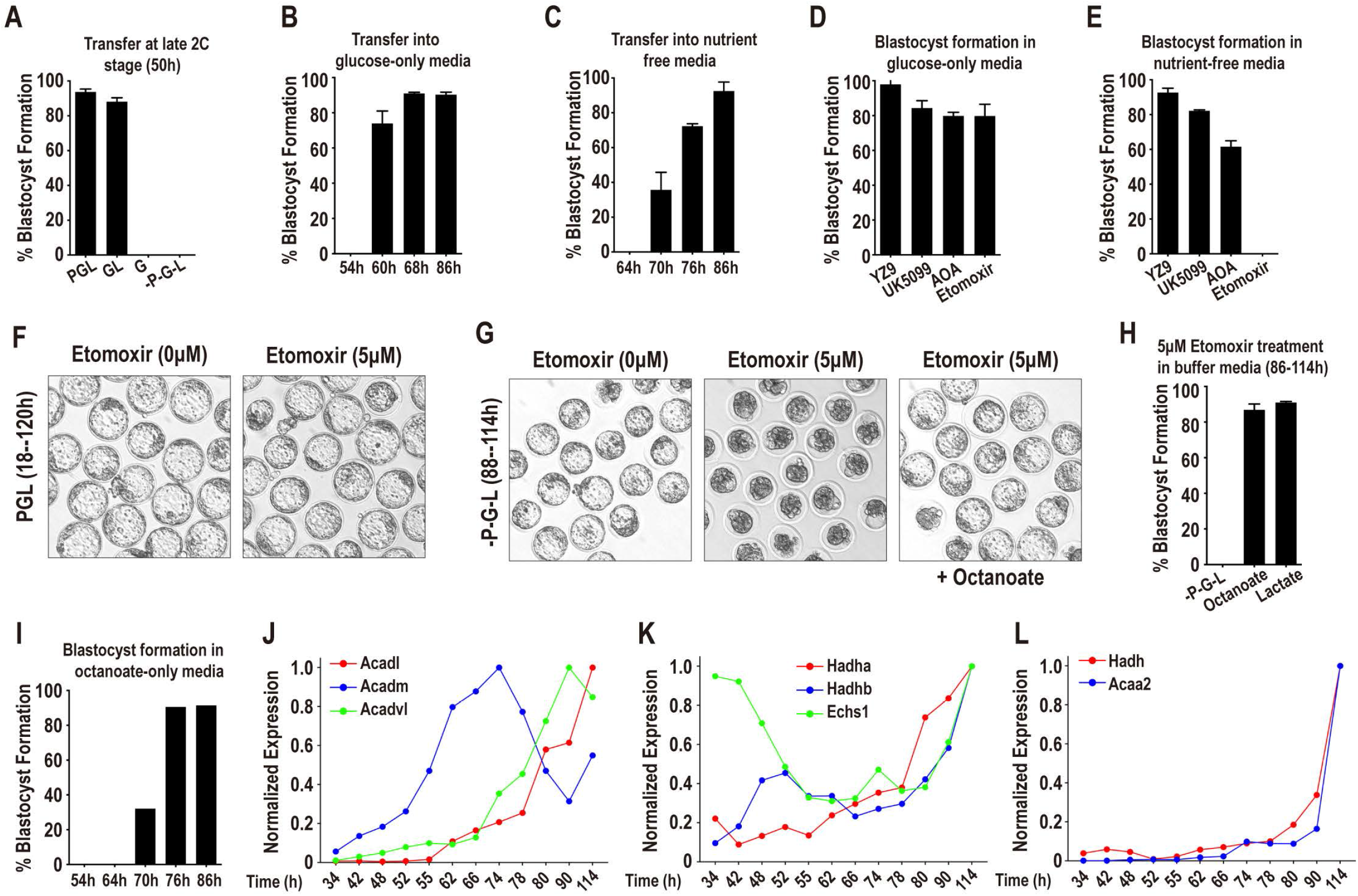
Further metabolic plasticity in later stage embryos. **(A**) At the late 2C stage (50h) pyruvate-deprived embryos require both glucose and lactate to form blastocysts. **(B)** Glucose alone, without pyruvate and lactate, can support blastocyst formation after the mid-4C stage (60h). **(C)** After 70h (compacting morulae) embryos form blastocysts in the absence of all three nutrients (-P-G-L). **(D)** Embryos transferred from PGL into glucose-only media at 86h are insensitive to inhibition of glycolysis (YZ9, 2μM), pyruvate entry into the mitochondria (UK5099, 1μM), the Mal-Asp shuttle (AOA, 0.5mM) and fatty acid oxidation (FAO) (etomoxir, 5μM). **(E)** In the absence of all nutrients the embryos remain insensitive to inhibition of glycolysis, pyruvate transport, and the Mal-Asp shuttle, but are highly sensitive to FAO inhibition (etomoxir) and are unable to form blastocysts. **(F)** Inhibition of fatty acid oxidation (FAO; etomoxir, 5μM) does not inhibit blastocyst formation in normal PGL media. **(G)** 200μM octanoate added to an otherwise nutrient free medium reverses the block in blastocyst formation caused by treatment with etomoxir. **(H)** Both octanoate (200μM) and lactate (2.5mM) compensate for FAO inhibition (etomoxir, 5μM). **(I)** Pre-8C embryos do not form blastocysts in nutrient-free media supplemented with 200μM octanoate. **(J-L)** Genes encoding enzymes of mitochondrial β-oxidation show high levels of expression in late morulae and blastocysts.

To determine how late stage (86h) embryos can possibly continue development either in the presence of glucose alone, or more so, how they continue development for up to 2 days beyond that without any exogenously provided nutrients, we used inhibitors of specific metabolic pathways that might be able to provide endogenous moieties to sustain development. We have shown that inhibition of the Mal-Asp shuttle, using AOA (Figure 4H) or glycolysis using YZ9 (Chi et al., 2020), under certain culture conditions, can block at earlier stages of development. In contrast, morulae (86h) developing in a medium that contains glucose, or one that is deprived of all nutrients are refractory to either AOA or YZ9 inhibition (Figure 5D, E).Thus, neither the Mal-Asp shuttle nor active glycolysis accounts for the embryo’s independence from external nutrients after the morula stage (Figure 5D, E) and we needed to look for alternative metabolic pathways.

To our surprise, embryos at this stage (86h) can develop into blastocysts even when the mitochondrial pyruvate carrier is inhibited by UK5099 (Figure 5D, E). Normally, pyruvate entry into the mitochondrion is a primary mechanism that is essential for the TCA cycle to generate energy. As a consequence, the development of zygotes (18h) or of 2C embryos (48h) is completely blocked by UK5099 even when they are cultured in all three nutrients (Nagaraj et al., 2017). However, later stage embryos (86h) do not require mitochondrial pyruvate entry to develop into fully expanded blastocysts with or without glucose (Figure 5D, E).

Finally, we investigated whether fatty acid oxidation is required under the nutrient-limiting conditions described above. The carnitine palmitoyl transferase (CPT) inhibitor etomoxir is highly effective in blocking the transfer of fatty acids into the mitochondrion where they can be used for β-oxidation. Embryos provided with pyruvate, lactate and glucose are insensitive to etomoxir, and develop into blastocysts even when treated from the zygote stage (Figure 5F). Morulae (86h) developing in glucose alone are also insensitive to inhibition by etomoxir and develop into blastocysts at the same frequency as untreated embryos (Figure 5D). In stark contrast, the embryos (86h) that are otherwise able to develop into blastocysts in a medium that lacks all nutrients, are entirely non-viable when treated with 5μM etomoxir (Figure 5E, G). This implies that fatty acid oxidation (FAO) plays a critical role in development during the external nutrient refractory stage. To test this model, we took advantage of the fact that medium chain fatty-acids, such as octanoate, do not require CPT to enter the mitochondrion. We find that added octanoate fully rescues the developmental block that is caused by simultaneous deprivation of all nutrients and treatment with etomoxir (Figure 5G, H). Importantly, the octanoate fueled capacity for the embryo to survive without added nutrients is a result of increased plasticity at later stages of preimplantation development since adding additional octanoate to a 2C/4C embryo, for example, cannot compensate for the lack of pyruvate, lactate and glucose (Figure 5I). Thus, it appears that the capacity of the embryo to utilize stored or provided lipids increases as development proceeds.

The important role of FAO during post-morula development led us to investigate the expression profile of genes related to mitochondrial β-oxidation in embryos grown in the control medium. The expression of the acyl-CoA dehydrogenase genes (*Acadm, Acadl, Acadvl*), which catalyze the first step in mitochondrial β-oxidation, is several-fold higher in late morulae and blastocysts than it is in the early cleavage stages (Figure 5J). The mitochondrial trifunctional enzyme components (*Hadha* and *Hadhb*) and crotonase (*Echs1*) increase in expression during the later stages of preimplantation development (Figure 5K). The most impressive increases are for short-chain (S)-3-hydroxyacyl-CoA dehydrogenase (*Hadh*) and medium-chain 3-ketoacyl-CoA thiolase (*Acaa2*), which are the final two components of the β-oxidation spiral (Figure 5L). Thus, all components of mitochondrial fatty acid β-oxidation increase substantially during the course of preimplantation development. The increased expression of the fatty acid oxidation enzymes during development reflects the metabolic reprogramming that facilitates the use of a wider range of resources at progressive stages of maturation. Their relative absence during the earliest stages of preimplantation renders the 2C embryos to be entirely reliant on pyruvate and lactate.

### Metabolic response to reductive stress

The gene expression data presented thus far reports the profile of preimplantation embryos cultured under normal (control) conditions. These data demonstrate dynamic expression patterns for a wide range of metabolic pathway genes. To determine if the transcriptome is sensitive to the nutrient content of the extracellular environment, we cultured embryos without pyruvate either from the zygote to the 2C stage (48h), or from the late 2C (50h) to the morula stage (78h). Overall, the expression of genes in mitochondrial fatty acid oxidation, purine and pyrimidine synthesis, the TCA cycle, glutamine metabolism, or glycine catabolism do not change as a group at either the 2C or morula stage following pyruvate withdrawal (Figure S5A-J). Furthermore, genes that encode enzymes that function in shuttles (such as the Mal-Asp and the glycerol-3P shuttle) that coordinate mitochondrial and cytoplasmic metabolism also do not increase when pyruvate is withdrawn. Rare exceptions to this invariance are seen for *Mdh1* (malate dehydrogenase), *Slc25a13* (mitochondrial aspartate and glutamate transporter) and *Slc25a1* (mitochondrial citrate transporter) which increase by 4-fold, 3-fold and 8-fold respectively, following pyruvate withdrawal (Figure S5B, F).

Glycolytic genes do not change in their relative expression level in 2C embryos that are cultured from the zygote stage without pyruvate. In stark contrast, we find that at the morula stage, pyruvate omission causes a coordinated increase in the expression of genes encoding glycolytic enzymes (Figure 6A). The largest increases are for *Gapdh* (6-fold), *Eno1* (7-fold), *Pkm* (8-fold), and *Ldha* (7-fold). Interestingly, when embryos are provided with α-KB and glucose (but not pyruvate and lactate), which allows progression past 2C, the glycolytic enzymes do not increase in expression in the morula (Figure S5J). To determine whether the increase in RNA levels upon pyruvate withdrawal leads to an increase in protein expression, we stained the embryos using specific antibodies for phosphofructokinase (PFK), glyceraldehyde-3-phosphate dehydrogenase (GAPDH) and pyruvate kinase (PKM2). We find that the protein levels of GAPDH, PFK, and PKM2 increase, with the rise in PKM2 especially marked (Figure 6B-C”, S5K-K”). Thus, following the 2C stage, the embryo responds to pyruvate withdrawal by mounting a coordinated transcriptional response that increases the abundance of glycolytic proteins.

**Figure 6.**
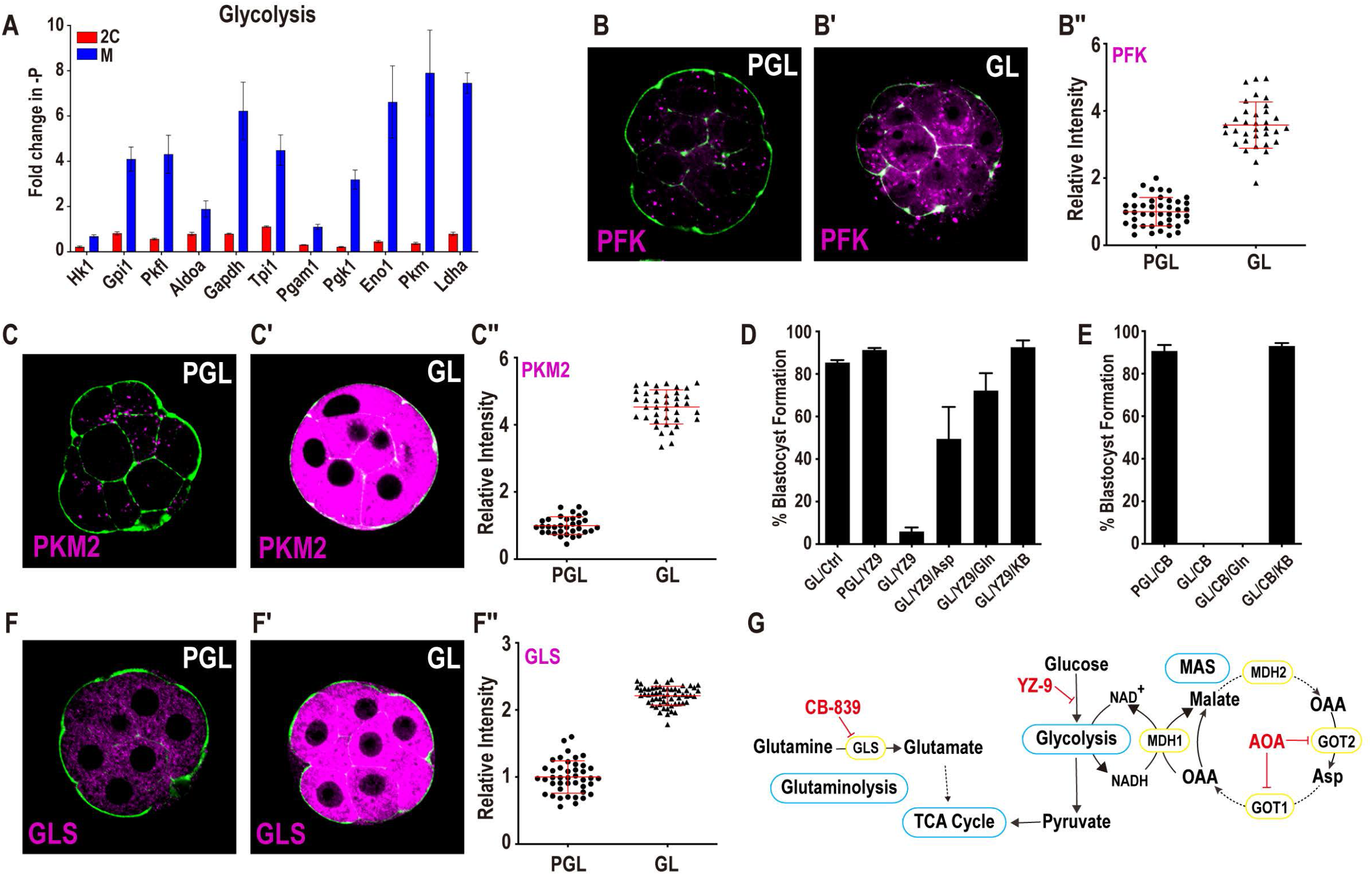
Metabolic response to reductive stress. **(A**) In morulae, but not in 2C embryos, the expression level of glycolytic genes increases when pyruvate is withdrawn. **(B-C”)** Upon pyruvate withdrawal, protein levels of the glycolytic enzymes PFK **(B-B”)** and PKM2 **(C-C”)** show significant increases. **(D)** The glycolysis inhibitor YZ-9 (2μM) blocks development of embryos deprived of pyruvate (GL) from the late 2C stage. This block is reversed by the addition of either Asp (1mM), Gln (1mM) or α-KB (1mM). **(E)** Glutaminase inhibition (200nM CB-839) does not block the development of embryos in PGL media. Embryos in GL media, however, block in development following CB-839 treatment. α-KB (1mM) suppresses the block, but Gln (1mM) does not. **(F-F”)** GLS protein increases when pyruvate is withheld. **(G)** Schematic diagram for glycolysis, the Mal-Asp shuttle, glutamine metabolism and their inhibitors.

In previous work we showed that inhibition of glycolysis (by either inhibiting PFK or PKM) blocks development when embryos are deprived of pyruvate, but not when pyruvate is included in the medium (Chi et al., 2020). We asked if the block in development of pyruvate deprived and glycolysis inhibited embryos will be rescued if the embryos are provided with an alternative means of generating NAD^+^. We achieve this by supplementation of the medium with either Asp, which generates NAD^+^ via the cytoplasmic arm of the Mal-Asp shuttle, or with α-KB, which generates NAD^+^ at the level of LDH (see Figure 6G). Both of these independent means of NAD^+^ augmentation rescue the block caused by simultaneous PFK inhibition and pyruvate withdrawal (Figure 6D). Thus, under stress conditions glucose metabolism, the Mal-Asp shuttle, and pyruvate/α-ketoacids are all able to generate NAD^+^ by parallel mechanisms.

In cancer cells and ESCs, a high level of glycolysis is often coupled with high rates of glutamine consumption. In the embryo, based on the RNA-seq analysis, this seemed unlikely as the transcript levels of *Gls*, *Gls2*, or *Glud1* remain unchanged upon pyruvate withdrawal (Figure S5H). However, a different conclusion arises when the glutaminase (GLS) protein, which converts glutamine to glutamate, is analyzed instead of its invariant RNA. GLS protein levels increase substantially upon pyruvate withdrawal (Figure 6F-F”) perhaps by a post-transcriptional mechanism similar to that seen in cancer models (Gao et al., 2009). Inhibition of glutaminase activity with the specific inhibitor CB-839 causes a block in development in pyruvate-deprived embryos but not when pyruvate is present (Figure 6E). The block in development caused by the glutaminase inhibitor is rescued when another α-ketoacid is included in the medium in place of pyruvate (Figure 6E). We conclude that the GLS protein level increases in response to reductive stress caused by pyruvate withdrawal, and thereby plays an active role in responding to a high NADH:NAD^+^ cellular environment.

### Mechanisms of metabolic reprogramming during reductive stress

Next, we investigated and characterized the mechanism by which embryos up-regulate glycolysis and glutaminase function in response to pyruvate withdrawal. Studies on other systems find that Myc and HIF-1α play important roles in glycolytic gene expression especially under various stress conditions (Gordan et al., 2007). Immunofluorescence analysis suggests that HIF-1α or HIF-2α are not stabilized in the embryo under any of the conditions that we use in this study (not shown). In contrast, nuclear Myc is detected in abundance in morulae that are deprived of pyruvate (Figure 7A-A”). Interestingly, by mechanisms as yet unclear, at the blastocyst stage, Myc expression is limited to the cells of the ICM (Figure 7B, (Claveria et al., 2013; Hashimoto and Sasaki, 2019)). Pyruvate withdrawal also leads to a robust increase in nuclearly-localized ICM specific Myc, while the TE cells continue to remain undetectable for Myc expression (Figure 7B-B”).

**Figure 7.**
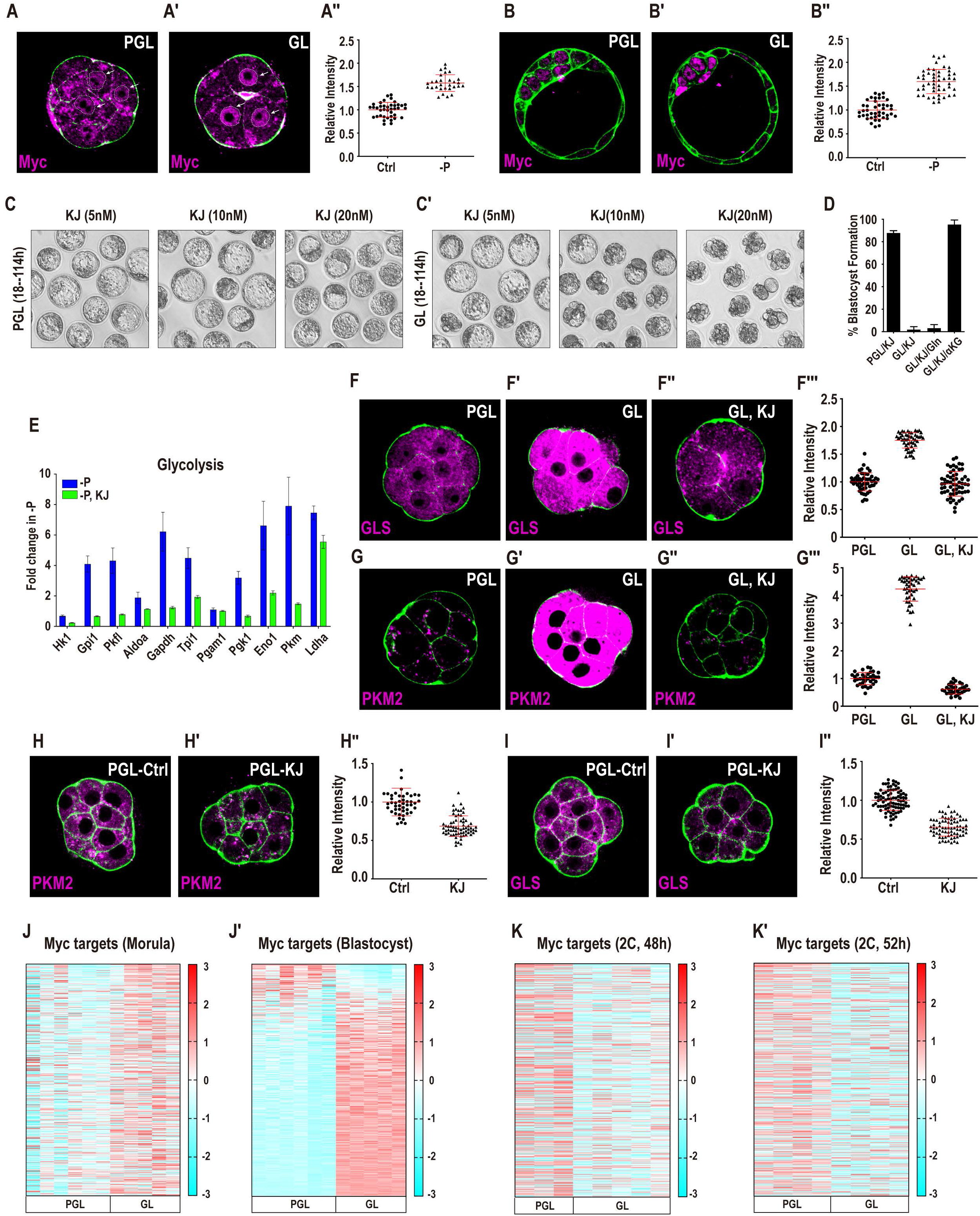
Mechanisms of metabolic reprogramming during reductive stress. **(A-B”**) Myc protein localization upon pyruvate withdrawal in the morula and blastocyst. **(A-A”)** In morulae, Myc is diffusely expressed in PGL **(A)**, but is nuclearly localized in GL **(A’)**. Quantitation in **(A”). (B-B”)** In blastocysts (114 h), Myc is expressed at low levels in PGL **(B)**. Pyruvate withdrawal causes a clear increase in Myc expression in the ICM but not in the TE **(B’)**. Quantitation in **(B”)**. **(C-C’)** Embryos cultured from the late 2C stage (50h) in GL **(C’)** but not in PGL **(C)** media block in development when treated with the Myc inhibitor KJ-Pyr-9 (abbreviated as KJ) at varying concentrations. **(D)** Embryos cultured in GL are sensitive to KJ (20nM). Gln (1mM) does not rescue, but supplementation with (1mM) α-KG (di-methyl) fully rescues the block. **(E)** Glycolytic genes do not increase in expression in pyruvate deprived embryos that are treated with 20nM KJ (green bars). The inhibitor-free sample (blue) is shared with Figure 6A, because these experiments were performed at the same time. **(F-G’”)** Myc inhibition blocks the increase in pyruvate sensitive genes GLS (**F-F’”**) and PKM2 (**G-G’”**). GLS **(F)** and PKM2 **(G)** levels rise in GL media **(F”, G’)**. KJ (20nM) prevents this increase in GLS **(F”)** and in PKM2 **(G”)**. Quantitation in **(F””, G’”)**. **(H-I”)** Myc functions under normal (PGL) growth conditions. Treatment with 20nM KJ (PGL medium, 2C, 50h) causes PKM2 **(H, H’)** and glutaminase **(I, I’)** protein levels to decrease. Quantitation in **(H’”, I”)**. **(J-K’)** A majority of Myc targets increase in expression at the morula stage **(J)** and more so in blastocysts **(J’)** following pyruvate withdrawal at the late 2C stage (50h). **(K, K’)** Myc targets do not increase in 2C embryos if they are cultured in a medium that lacks pyruvate from the zygote stage.

We inhibited the activity of Myc using a specific inhibitor (KJ-Pyr-9, which prevents Myc-Max interaction), in order to assess the functional role of Myc in protecting the embryo during reductive stress. Embryos provided with pyruvate are resistant to treatment with 20nM KJ-Pyr-9 and readily form blastocysts under these conditions (Figure 7C, D). In contrast, without access to pyruvate, embryos are highly sensitive to Myc inhibition, and exhibit a complete developmental block prior to the morula stage (Figure 7C’ D). This Myc inhibition also abrogates the increases in glycolytic gene expression and the rise in PKM2 and GLS proteins that is observed following pyruvate withdrawal (Figure 7E-G’”). There is not a significant block in morula formation when Myc is inhibited under normal culture conditions. However, the protein levels of PKM2 and GLS are significantly reduced upon Myc inhibition even with pyruvate, lactate and glucose present (Figure 7H-I”) suggesting a measure of metabolic control by Myc during normal development that becomes far more prominent under conditions of reductive stress.

The above results are substantiated by RNA-seq experiments, which reveal a dramatic upregulation of Myc targets at both the morula and blastocyst stages upon pyruvate withdrawal (Figure 7J, J’). We used two independently derived Myc target gene-lists; (Morrish and Hockenbery, 2014) - list 1 and (Kim et al., 2008) - list 2. For either list, 90% of the Myc targets are affected by pyruvate withdrawal (Figure 7J, list 1 and Figure S6B, list 2). Importantly, there is no increase in Myc target genes when embryos are cultured with glucose as the sole nutrient source (with added α-KB) (Figure S6C, C’). We also find that Myc target genes do not increase in expression in 2C embryos when pyruvate is withheld from the zygote stage (Figure 7K, K’). This result is reasonable since transcription of zygotic genes does not take place under these conditions. This would, in part, explain why early 2C embryos are far more sensitive to pyruvate withdrawal when compared with embryos at later stages of development. Increased expression of Myc targets upon reductive stress is only possible in embryos that have completed ZGA and before that event, the embryo is unable to respond to stress through the coordinated upregulation of the Myc-related network.

## Discussion

Unique features of the preimplantation steps of mouse embryonic development are its minimal level of interactions with the environment, with no requirement for cytokine signals; small maternal stores in the form of yolk; low bioenergetic activity; and a reliance on specific nutrients. Preimplantation developmental steps follow a set program that is largely driven by metabolic processes. The earliest cleavage is the most rigid, controlled by the pyruvate/lactate system and a disproportionately large maternal deposit of the LDHB protein that maintains high NADH and enforces a pyruvate dependent initiation of ZGA. These metabolic requirements are relaxed at the later steps with an expansion of metabolic plasticity. This expansion initiates as maternal LDHB levels are lost and are not replenished by the embryo, allowing several metabolic shuttles to operate that rebalance the NADH/NAD^+^ ratio. This also enhances the contribution of glycolysis and equilibrates the TCA cycle. The repertoire of nutrients that can sustain these developmental steps expands such that, at the late morula stage, embryos can proceed to the expanded blastocyst with no requirement for any externally added nutrient. The gradual expansion in metabolic plasticity is essential for development to proceed from the zygote to the expanded blastocyst. The redox state of the embryo plays a critical role in this process since it can alter the outcome of enzymatic processes, and it directs a highly choreographed metabolic sequence of enzyme deposition, degradation, their networked transcriptional regulation and function that provides the proper set of metabolites, bioenergetics, biosynthesis and redox balance. The metabolic strategy employed by the embryo in its nutrient utilization is in many ways distinct to that followed by cancer cells. However, we find that stress conditions cause the embryos to adapt in ways very similar to that of cancer cells and cultured ESCs. Thus, expansion of metabolic plasticity is a two-edged sword used for normal development but also contributing to uncontrolled growth when expanded in abnormal genetic backgrounds evident in disorders such cancer.

There are a number of related factors that generate the constraints which restrict the nutrients that can sustain early stage embryos and that bring about increased metabolic plasticity later in development. For example, zygotes and early 2C embryos provided with lactate alone are unable, following uptake, to intracellularly convert enough of this lactate to pyruvate for it to be used as a nutrient. However, when pyruvate (or another α-ketoacid) is provided along with lactate in the medium, now following uptake, further conversion of intracellular lactate to pyruvate is permitted. This is because the pyruvate facilitates regeneration of NAD^+^. As development proceeds beyond the 2C stage, components of the Mal-Asp shuttle, the glycerol-3P shuttle and the mitochondrial electron transport chain all increase in abundance, and consequently embryos are able to regenerate NAD^+^ that is consumed during lactate to pyruvate conversion, and are able to develop using lactate without added pyruvate.

Glucose cannot support development until the 8C stage, and when both pyruvate and lactate are absent, early embryos are not viable. In our model, this is because of the extraordinarily high LDHB activity in the embryo before this stage, relative to glycolysis. When glucose is provided to embryos that are deprived of both pyruvate and lactate, only a small amount of pyruvate results from glycolysis and is virtually all converted to lactate. At this stage, embryos cannot regenerate NAD^+^ at sufficient rates to convert lactate back to pyruvate before the lactate is exported from the cell. However, glucose can serve as an effective nutrient in the absence of pyruvate and lactate when supplemented with an α-ketoacid. These compounds work to restore NAD^+^ levels thus disfavoring the conversion of glycolysis generated pyruvate into lactate. Increasing NAD^+^ levels may also increase the activity of GAPDH, a key glycolytic enzyme that is inhibited when NAD^+^ is low. We conclude that the resulting redox shift mitigates the impact of having such high levels of LDHB present. Thus, large amounts of LDHB and an inability to regenerate cytoplasmic NAD^+^ renders early mouse embryos unable to survive using glucose as a nutrient.

As morulae form, the level of LDHB continues to decrease, whereas the expression of the shuttle components increases, as does the rate of mitochondrial oxygen consumption. Consequently, at the early morula stage, glucose (without added α-ketoacids) is able to support development even when both pyruvate and lactate are absent. The ability of embryos to develop without pyruvate and lactate is not due to a switch to a glucose-based metabolism that occurs during normal development, but rather represents an increase in plasticity that is caused by a decrease in LDHB levels and an increased ability of the embryo to regenerate NAD^+^.

Following compaction, metabolic plasticity increases further, and embryos are now able develop into expanded blastocysts even when no external nutrients are provided. We discovered a critical role of fatty acid oxidation (FAO) under these conditions. Interestingly, the requirement for FAO is not a critical factor in development when pyruvate, lactate or glucose are present. Thus, the ability to choose between pathways based on nutrient availability is an additional mechanism for metabolic plasticity. *In utero*, at the late blastocyst stage, the embryo is known to move through a nutrient depleted environment and therefore, this requirement for FAO is likely invoked during normal *in vivo* conditions. This assumption is consistent with our gene expression data that found that multiple mitochondrial fatty-acid oxidation components are expressed at the highest levels in the morula and blastocyst. Thus, to the growing list of mechanisms underlying increasing metabolic plasticity with development, we can add switching on of catabolic pathways that are not present, or are inactive, during the earliest stages of development.

In general, the embryos maintain a “thrifty” metabolic profile, storing several maternal products for later stages and utilizing limited environmental resources at the earliest stages as a survival strategy. In this sense the embryo is quite different from cancer cells and ESCs. Consequently, for the most part, the metabolic characteristics of the embryo contrast markedly with those of cultured embryonic stem cells or cancer cells, which avidly take-up glucose and glutamine and do not require either pyruvate or lactate (but see (Faubert et al., 2017; Hui et al., 2017)). There are important similarities, however, that arise under conditions of stress.

The embryo is also equipped to diverge from its “thrifty” metabolic state in response to changes in the extracellular environment, particularly those that cause redox stress. A major adaptive response to stress, we find, involves a c-Myc dependent process. Once again, Myc is a known player in cancer metabolism (Dang, 2012), and its adaptive role in developmental metabolism is both important and interesting given the large number of differences that are apparent between the two systems. And yet, as our studies with Myc highlight, the components of the metabolic networks in embryos under stress and cancer cells are the same, and analysis of the developmental system provides numerous parallels in how embryos and cancer cells adapt to changing metabolic needs. It seems that stress situations bring out similarities between developmental and cancer systems.

In addition to increasing plasticity, the metabolism of the preimplantation embryo also shows a remarkable level of self-sufficiency, and a very limited reliance on its environment. For instance, pyruvate/lactate can support development of the embryo until the compacted morula stage without the provision of glucose, amino acids, nucleotide precursors, vitamins, or any form of lipid. This self-sufficiency extends beyond nutrient uptake, as the embryo is able to develop without the input of growth factors, cytokines or any other form of external signaling molecules. Previously we provided evidence that metabolism plays a critical role in developmental decisionmaking that would, in other circumstances, be achieved by a cell’s extensive reliance on external signaling input. For instance, we have shown previously that pyruvate is critical for ZGA, and that glucose metabolism controls TE cell fate determination, distinguishing them from cells of the ICM. Thus, in the embryo, the metabolic transitions that facilitate increased plasticity are not providers of passive support to the core events of the developmental program, but are, in fact, central players that inform developmental processes.

Following ZGA, the transcriptional profile of genes related to metabolic pathways changes in a concerted fashion based on a pre-programmed cascade of transcriptional events. The increased expression of these newly formed metabolic enzymes (that favor plasticity) contrasts with the decrease in the maternally deposited pool (that tend to favor rigidity). The rate of change in the levels of these two sets of metabolic enzymes sets a developmental timer, and only once a high ratio of zygotic to maternal enzymes is reached can the embryo develop into diverse cell types, such as ICM or TE cells. The inflexible metabolism of the early embryo favors the storage of nutrients for later stages of development. Under conditions of stress, the early embryo achieves greater plasticity by reprogramming its metabolism. However, metabolic reprogramming in response to stress will deplete nutrient stores, and thus impact later stages of development. In future studies, it will be important to determine if changes in metabolic plasticity, stress-induced or not, is one of the causes of early loss of pregnancy or results in birth defects.

## Author Contributions

M.S.S., F.C. and U.B. conceived the project, wrote and discussed the manuscript. M.S.S, and F.C. performed experiments. U.B. secured funding and provided mentorship.

## Acknowledgments

We thank Raghavendra Nagaraj and all members of our laboratory for their suggestions. We thank Daniel Braas and Johanna ten Hoeve-Scott at the UCLA Metabolomics Center for their generous help with metabolomics analysis. Sequencing was performed by Suhua Feng of the BSCRC Sequencing Core, and we thank Steve Jacobsen for his support with sequencing. We appreciate many inputs by Tom Graeber, Heather Christofk and Hilary Coller. Christofk and Coller also helped by providing many antibody reagents.

This work was supported by the NIH Director’s Pioneer Award to U.B. (DP1DK098059). U.B. is supported by the NCI grant R01 CA217608, NHLBI grant R01 HL067395. F.C. is supported by a China Scholarship Council Award, a California Institute for Regenerative Medicine Predoctoral Fellowship, and the MBI Whitcome Predoctoral Fellowship. M.S.S was supported by the CTSI Iris Cantor Women’s Health Center Award. Finally, we are grateful to Owen Witte and the Broad Stem Cell Research Center for several innovation and research awards to UB and the BSCRC predoctoral fellowship support to F.C. We are particularly grateful to the Gillian S. Fuller Foundation for their support of BSCRC in general and this work in particular.

## Experimental Model and Subject Details

### Lead Contact and Materials Availability

Further information and requests for resources and reagents should be directed to and will be fulfilled by the lead contacts, Mark Sharpley or Utpal Banerjee

### Mouse embryo culture

All animal care and procedures used in this study are approved by the Animal Regulatory Committee (ARC) of the University of California at Los Angeles (UCLA).

Mouse zygotes and preimplantation embryos were collected from super-ovulated 4-week old C57BL/6J X C3He (Jackson Labs) F1 females. Mice were super-ovulated by peritoneal injection of 7.5 IU of PMSG (Pregnant Mare Serum Gonadotropin) to stimulate egg production, followed by 7.5 IU of hCG (human Chorionic Gonadotropin) 48h after PMSG. Embryos were obtained by mating the super-ovulated females with C57BL/6 X C3He F1 males. Mating was confirmed by the presence of the vaginal plug. For isolation of fertilized 1-cell zygotes, super-ovulated females were euthanized 18h post hCG and zygotes were dissected out of the ampulla in the oviduct. The embryo cumulus complexes were treated with 300μg/ml of hyaluronidase to disperse the cumulus cells, washed in mKSOM medium without pyruvate/glucose and transferred to the appropriate culture medium and cultured at 37°C in 5% CO_2_. All mouse embryos used in this study were cultured in a modified KSOM medium whose composition is identical to KSOM in salts, glucose, lactate and pyruvate (95mM NaCl, 2.5mM KCl, 0.35mM KH_2_PO_4_, 0.20mM MgSO_4_, 25mM NaHCO_3_, 1.71mM CaCl_2_ 0.01 mM EDTA, 0.20mM glucose, 5mM L-lactate, 0.20mM pyruvate) but was devoid of all amino acids and BSA. The medium also contained 0.01% PVA (polyvinyl alcohol).

### Method Details

#### Reagents

The following antibodies and drugs were used in this study. Fluorescein Phalloidin (Thermo Fisher #F432, at 1:2000 dilution), rabbit anti-PKM2 (CST #4053), goat anti-PFK1 (Santa Cruz #sc-31712), mouse anti-GAPDH (Santa Cruz #sc-32233), rabbit anti-GLS (Abcam #ab93434), rabbit anti-GLUD1 (Abcam #ab168352), and rabbit anti-c-Myc (Santa Cruz #sc-764). All the antibodies are used at 1:100 dilution for IF.

The chemical reagents used in this study were: YZ9 (Cayman Chemical #15352, 1μM), KJ-Pyr-9 (Cayman Chemical #19116, 20nM), CB-839 (Cayman Chemical #22038, 100nM), O- (Carboxymethyl)hydroxylamine hemihydrochloride (AOA) (Sigma #C13408, 0.5mM), L-Methionine sulfoximine (MSO) (Sigma #M5379, 1mM), Etomoxir (Sigma #E1905, 5μM), UK-5099 (Sigma # PZ0160, 500nM). The inhibitors were used at the time indicated in the text to treat the embryos. To avoid absorption of inhibitors into the oil-overlay, hydrophobic inhibitors were provided to embryos that were cultured under oil-free conditions.

#### Antibody staining for immunofluorescence

Embryos were fixed in 4% paraformaldehyde for 30 min at room temperature, permeabilized for 30 min in PBS with 0.4% Triton (PBST), blocked in PBST with 3% albumin (PBSTA) for 30 min and incubated with the desired primary antibody in PBSTA overnight at 4°C. The following day the embryos were washed in PBST 4 times for 10 min each, blocked with PBSTA, incubated with the appropriate secondary antibody (1:500 dilution) and DAPI overnight at 4°C. Embryos were washed again 3 times for 10 min each in PBST, deposited on glass slides and mounted in Vectashield (Vector Laboratories) medium. Images were captured using Zeiss LSM700 or LSM880 confocal microscopes.

#### RNA sequencing experiments

The RNA-seq libraries were generated from early mouse embryos using the Smart-seq2 protocol as published with minor modification (Picelli et al., 2014). Cells were lysed in 0.1% (vol/vol) Triton X-100 lysis buffer containing 2.5μM of oligo-dT30VN primer, 2.5mM of dNTP mix and 1U/μl of RNase inhibitor. After 3 mins lysis at 72°C, the Smart-seq2 reverse transcription mixture was added for reverse transcription. After pre-amplification and AMPure XP beads purification, 0.5ng of cDNA was used for the Tn5 tagmentation reaction. Libraries were sequenced on Illumina HiSeq-4000 according to the manufacturer’s instruction. The RNA sequencing data were analyzed using Partek software suite. The reads data were aligned to mouse reference genome mm10-ensembl transcripts (release 97) using the STAR package that is embedded in Partek.

#### Measurement of metabolite levels

The metabolites are measured using the procedure described previously (Nagaraj et al., 2017) (Sullivan et al., 2018). For samples analyzed under different conditions, embryos were very briefly washed in ice-cold 150mM ammonium acetate, and transferred into 80% methanol. Dried metabolites were re-suspended in 20μl 50% ACN and 5μl was injected for chromatographic separation on the UltiMate 3000 RSLC (Thermo Scientific) UHPLC system which is coupled to a Thermo Scientific Q Exactive that is run in polarity switching mode. For the UHPLC separation, the mobile phase comprised (A) 5mM NH_4_AcO, pH 9.9, and (B) ACN. A Luna 3mm NH2 100A (150 × 2.0mm) (Phenomenex) column was used with a 300 μL/min flow-rate. The gradient ran from 15% A to 95% A in 18 min, followed by an isocratic step for 9 min and re-equilibration for 7 min. The remainder of the sample was diluted 2.67-fold and 10μl were applied to a Thermo Scientific Ion Chromatography System (ICS) 5000 that is also coupled to a Thermo Scientific Q Exactive run in negative polarity mode. The gradient ran from 10mM to 90mM KOH over 25 min with a flow rate of 350μl/min. The settings for the HESI-II source were: S-lens 50, Sheath Gas 18, Aux Gas 4, spray heater 320°C, and spray voltage −3.2 kV. Metabolites were identified based on accurate mass (±3 ppm) and retention times of pure standards. Relative amounts of metabolites and contribution of ^13^C labeled nutrients (following correction for natural abundance) were quantified using TraceFinder 3.3. The data were analyzed using the Prism software package.

NAD^+^ and NADH levels were determined using the NAD^+^/NADH Glo cycling assay, according to the manufacturer’s instructions (Promega, WI, USA). Briefly, both NAD^+^ and NADH were extracted from the embryos in a carbonate buffered base solution containing 1% DTAB, 10 mM nicotinamide, 0.05% Triton X-100, pH 10.7. The embryos were freeze-thawed in liquid nitrogen and centrifuged through a filter with a 10kDa mW cut-off. Samples for NAD^+^ determination were neutralized with 0.4 N HCl. Both NAD^+^ and NADH samples were heated for 20 min at 60°C. The assay reagent was added to the samples, which were then incubated for 3h for cycling, after which the luminescence was determined.

#### Quantification and Statistical Analysis

Statistical parameters are reported in the figures and figure legends. Data is considered significant if p < 0.05. Statistical analysis was performed using GraphPad Prism software.

## Supplementary Figure legends

**Figure S1, related to Figure 1:**
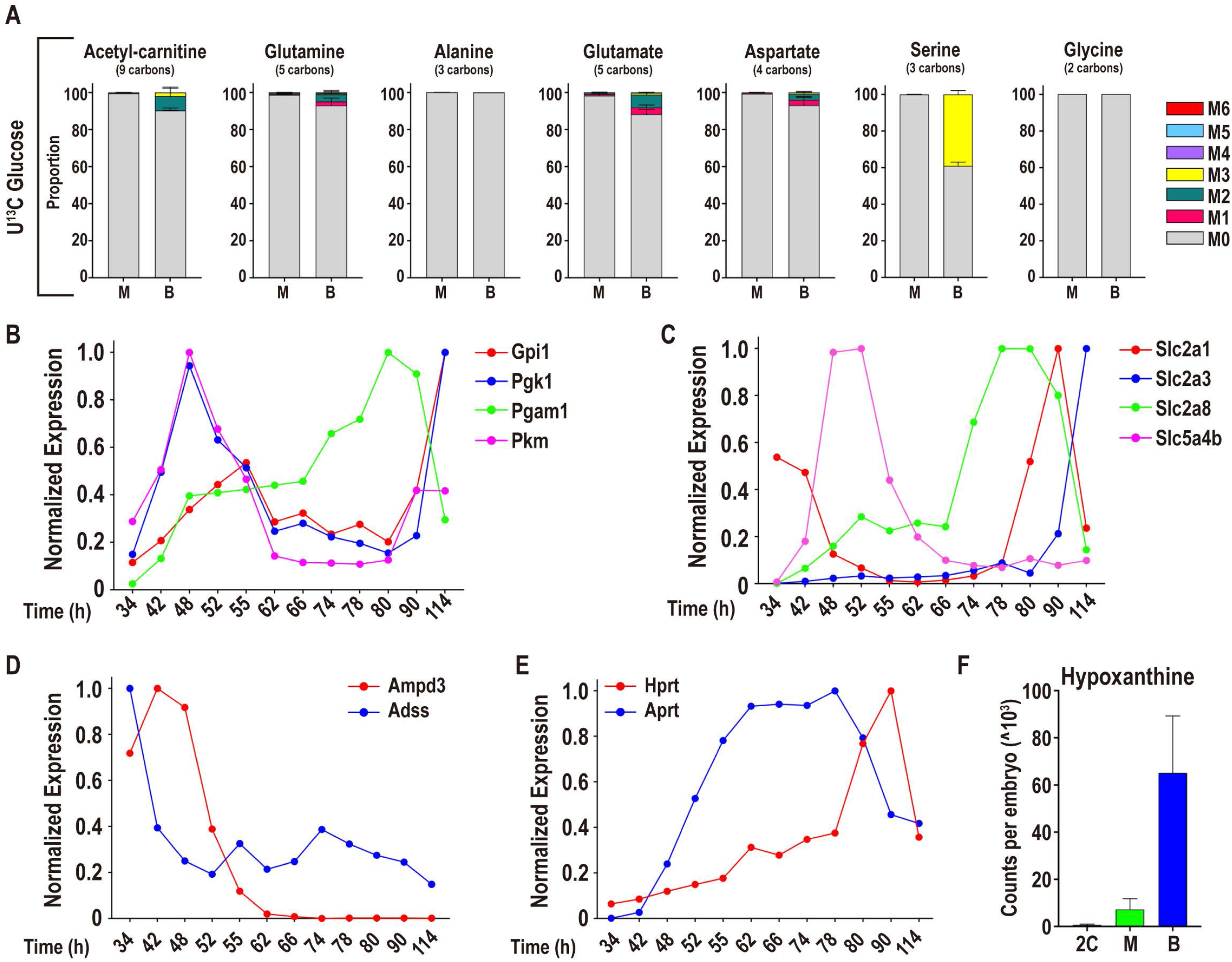
Glucose metabolism during preimplantation development. **(A)** U-^13^C glucose only contributes minor amounts of carbon to acetyl-carnitine and amino acids, with a measurable contribution only at the blastocyst stage. At the blastocyst stage, serine is partially labeled by glucose, but glucose does not contribute to glycine formation. Gray: M0 unlabeled; colors: labeled as marked. Note that the “fully labeled” color depends on the total carbons in the metabolite (e.g. fully labeled is M5 for Glu and M4 for Asp). **(B)** Expression of genes encoding glycolytic enzymes between early 2C (34h) and fully expanded blastocyst (114h) stages. A subset of glycolytic genes peak in gene expression at the 2C stage, and then increase during blastocyst formation. See also Figure 1E, F. **(C)** The glucose transporters *Slc2a1* (*Glut1*) and *Slc2a3* (*Glut3*) peak in gene expression at the blastocyst stage. *Slc2a2* (*Glut2*) and *Slc2a4* (*Glut4*) are not expressed in preimplantation embryos at significant amounts. Slc2a8 peaks in expression during the morula stage, though the function of this transporter is unclear. The sodium coupled glucose transporters *Sglt1* (*Slc5a1*) and *Sglt2* (*Slc5a2*) are not expressed. A third sodium coupled glucose transporter *Sglt3* (*Slc5a4b*) is expressed in 2C embryos. *Slc5a4b* may function primarily as a glucose sensor rather than a transporter. **(D)** The expression of genes that encode the purine nucleotide cycle enzymes *Ampd3* and *Adss* is greatest during the earliest stages of development. **(E**) The expression of genes that encode the purine salvage enzymes *Hprt* and *Aprt* peaks during the morula and blastocyst stages. **(F)** The amount of hypoxanthine, a substrate for the salvage enzyme HPRT, that is present in the embryo increases dramatically between the 2C stage and blastocysts.

**Figure S2, related to Figure 2:**
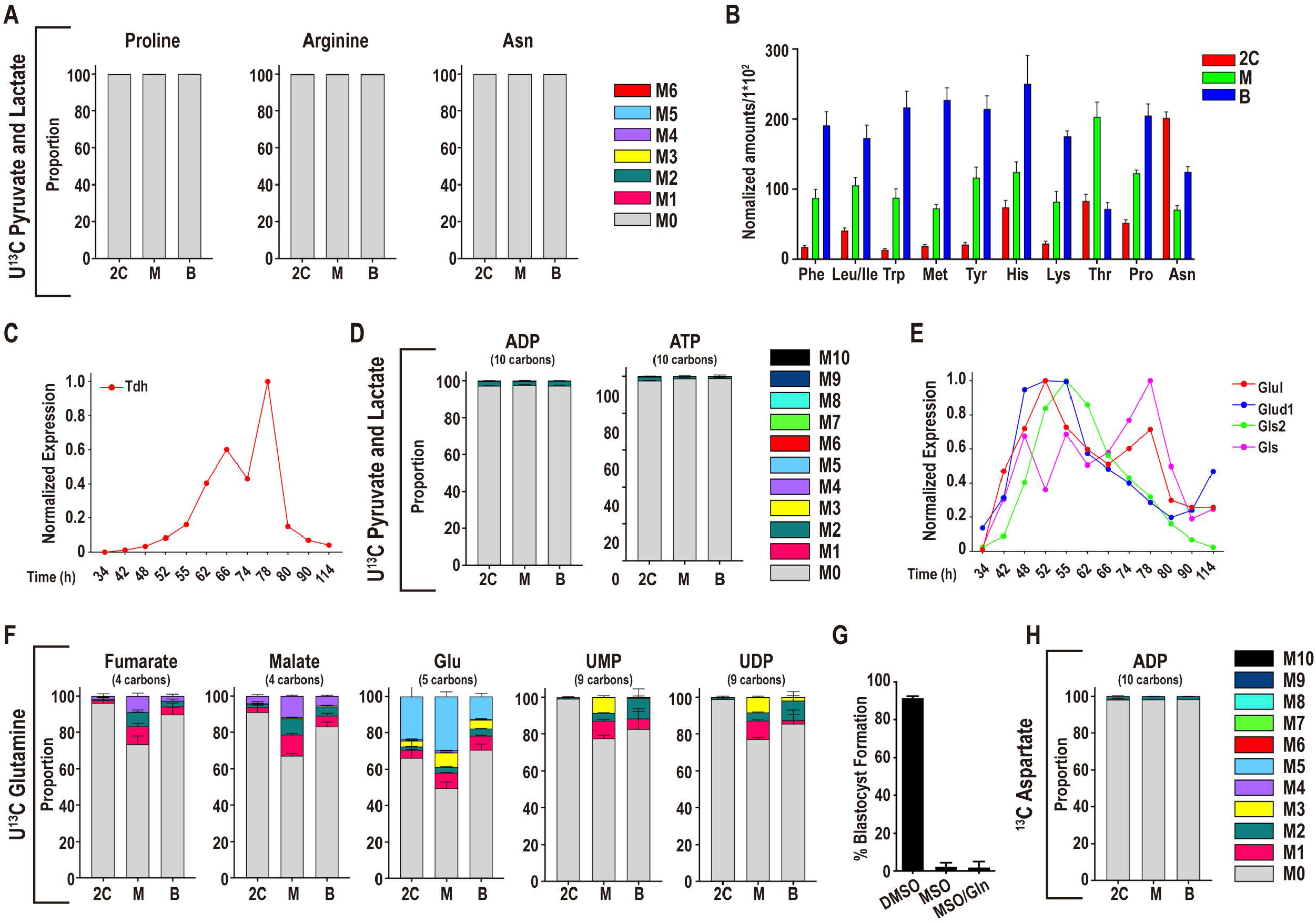
Pyruvate and lactate metabolism during preimplantation development. **(A)** U-^13^C pyruvate and U-^13^C lactate do not contribute to the amino acids, proline, arginine, and asparagine. In all figure panels, stages are represented as 2C: 2-cell, M: morula, B: blastocyst. **(B**) Scaled amounts of amino acids at each stage. **(C)** The expression of threonine dehydrogenase (*Tdh*) is greatest at the morula stage **(D)** U-^13^C pyruvate and U-^13^C lactate do not contribute carbons to purine nucleotide synthesis. **(E)** The genes that encode enzymes that function in glutamine metabolism exhibit the highest expression during the 4C and 8C stage. **(F)** U-^13^C Gln only contributes minor amounts of carbon to the TCA cycle, amino acids and pyrimidine nucleotides. **(G)** The glutamine synthetase inhibitor MSO blocks development when added to late 2C stage embryos (50h). Provision of exogenous glutamine does not rescue the developmental block that is caused by MSO. **(H)** U-^13^C aspartate does not contribute carbons to purine nucleotides.

**Figure S3, related to Figure 3:**
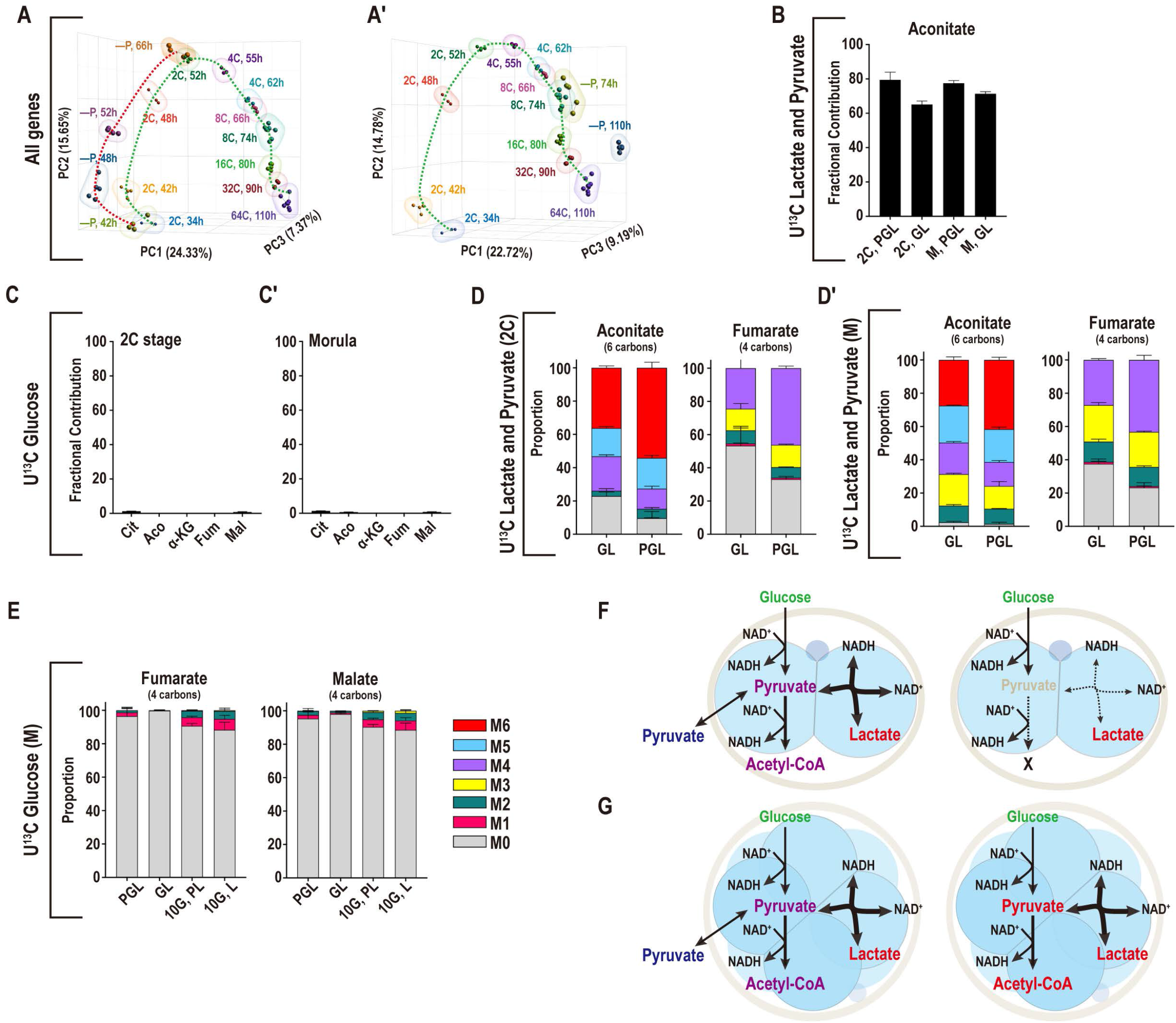
Adaptation to nutrient conditions. **(A, A’)** PCA analysis of the transcriptome (34112 genes) of embryos developing with and without pyruvate. Pyruvate is removed from the zygote stage **(A)** or the late 2C stage **(A’)**. **(B)** In both 2C embryos and morulae, U-^13^C pyruvate and U-^13^C lactate contribute a majority of carbons to the total aconitate pool in the presence (PGL) or absence (GL) of pyruvate. **(C, C’)**. Contribution of glucose to the total carbon pool of TCA cycle metabolites in embryos that are deprived of pyruvate. In both 2C embryos **(C)** and morulae **(C’)**, U-^13^C glucose does not contribute carbons to the TCA cycle when pyruvate is withheld. **(D, D’)** At both the 2C **(D)** and morula **(D’)** stages, the isotopologues of aconitate and fumarate that are formed from U-^13^C lactate alone (no pyruvate is present, GL) are similar to the isotopologues that are formed when both U-^13^C pyruvate and U-^13^C lactate (PGL) are present. **(E)** The contribution of U-^13^C glucose to the TCA cycle metabolites malate and fumarate remains very low in morulae in GL media (lacking pyruvate), even when a high amount of glucose (10mM instead of the normal 0.2mM) is provided (10G, L). **(F, G)** Schematic illustrating pyruvate metabolism in **(F)** 2C embryos, and **(G)** morula. At both stages, when pyruvate and lactate are present, lactate and pyruvate contribute to the TCA cycle, and glucose does not. **(F)** In 2C embryos, when pyruvate is absent, neither glucose or lactate are able to support the maintenance of TCA cycle metabolite pools, whereas in morulae **(G)** when pyruvate is absent, lactate, but not glucose, supports the maintenance of the TCA cycle metabolite pools.

**Figure S4, related to Figure 4:**
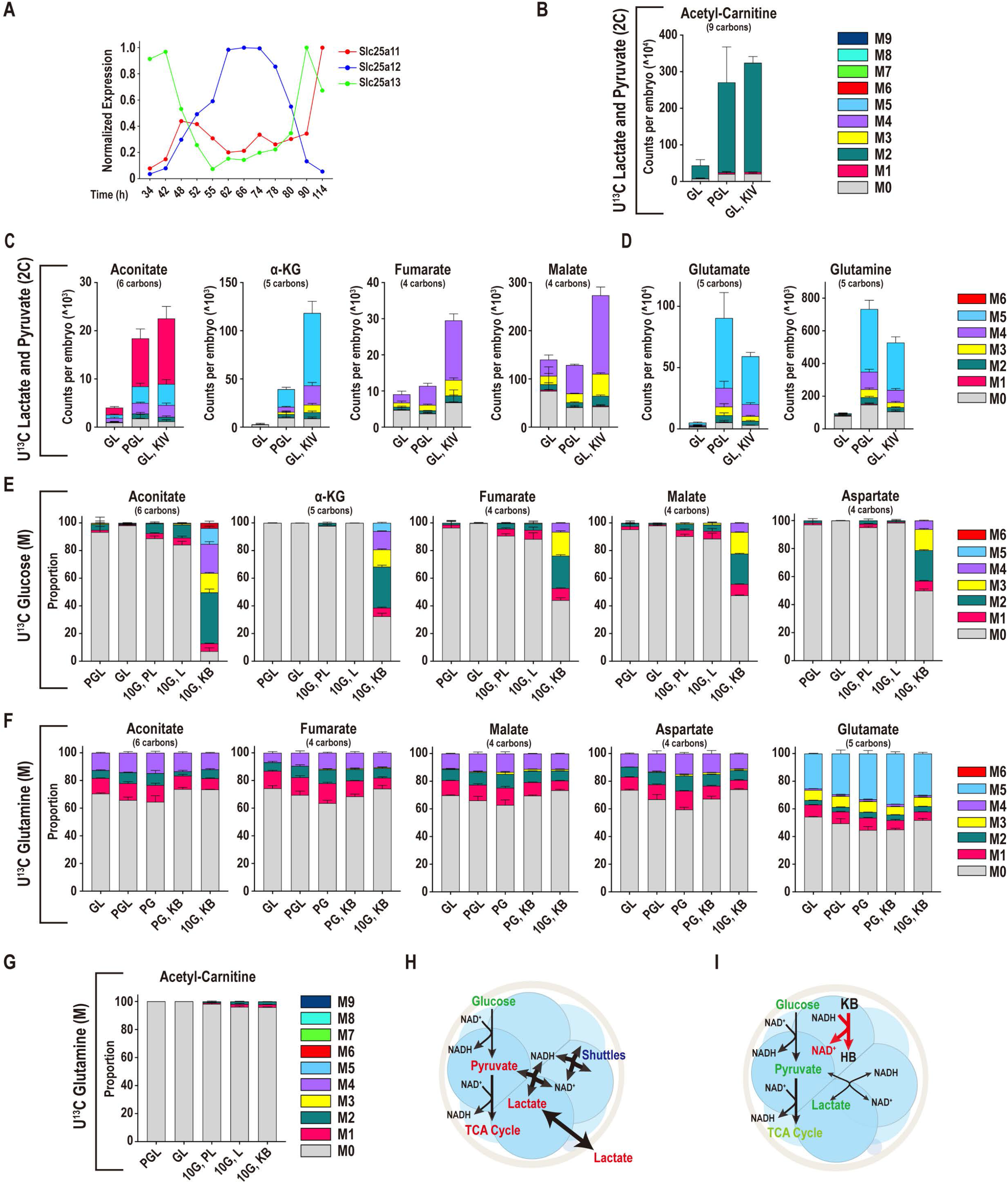
Mechanisms of metabolic plasticity. **(A)** Expression of the malate-aspartate shuttle components that transport aspartate and glutamate (*Slc25a12, Slc25a13*) or malate and α-KG (*Slc25a11*) across the mitochondrial membrane. **(B-D)**. Rescue of metabolites by α-KIV in pyruvate-deprived embryos. U-^12^C α-KIV restores acetyl-carnitine **(B)**, TCA cycle metabolites **(C)**, and Gln and Glu **(D)** levels in 2C embryos that are deprived of pyruvate. U-^13^C lactate contribution to the TCA cycle metabolites is similar in PGL and in KIV supplemented embryos in GL. The PGL and GL control samples are shared with Figure **3F**, because these experiments were performed at the same time. **(E)** U-^13^C glucose (0.2 mM in PGL/GL, or 10mM 10G, PL/L/KB) contributes insignificant amounts of carbon to TCA cycle metabolites and Asp in morulae in either PGL or GL media. In contrast, glucose contributes substantially when both pyruvate and lactate are omitted if α-KB is provided. The control samples are shared with Figure **3E**, because these experiments were performed at the same time. **(F, G)** Contribution of U-^13^C Gln to the TCA cycle, amino acids or acetyl-carnitine when both pyruvate and lactate are absent. U-^13^C glutamine contributes minor amounts of carbon to TCA cycle metabolites and amino acids **(F)** and acetyl-carnitine **(G)** even when both pyruvate and lactate are omitted from the medium and unlabeled 10mM glucose and α-KB are provided. **(H, I)** Schematics illustrating how **(H)** morula stage embryos use lactate to develop in the absence of pyruvate, and **(I)** depicting how the presence of α-KB facilitates the use of glucose as a source of carbon for the TCA cycle.

**Figure S5, related to Figure 6:**
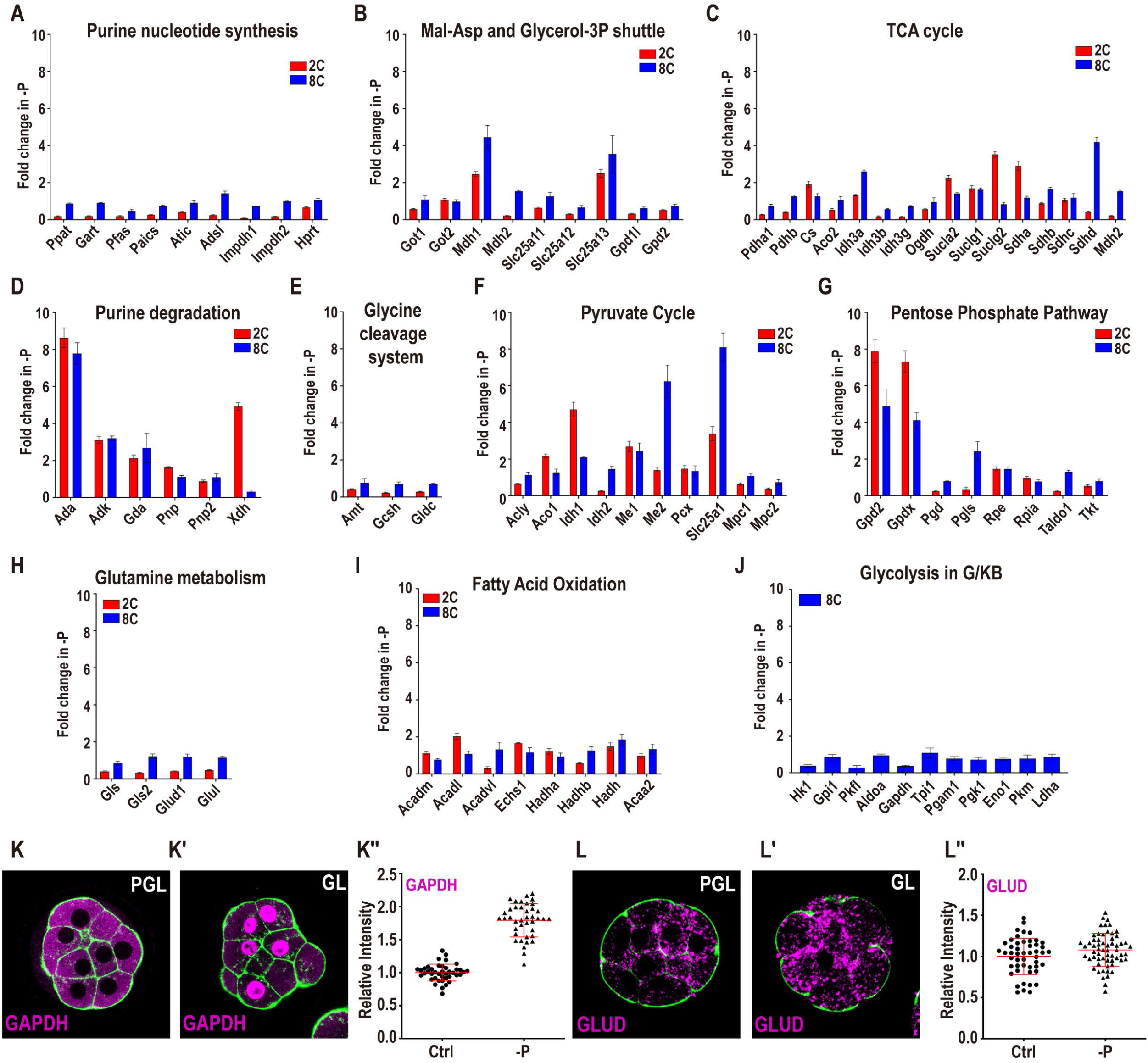
Metabolic response to reductive stress. **(A-I)** Fold change in expression of genes functioning in metabolic pathways in 2C embryos following pyruvate withdrawal from the zygote stage (red bars), and morulae, following pyruvate withdrawal at the late 2C stage (blue bars). **(A)** purine nucleotide synthesis, **(B)** the Mal-Asp shuttle and glycerol-3P shuttle, **(C)** TCA cycle, **(D)** purine degradation, **(E)** the glycine cleavage system, **(F)** the pyruvate cycle, **(G)** the pentose phosphate pathway, **(H)** glutamine metabolism, and **(I)** mitochondrial fatty acid oxidation. **(J)** There is not an increase in glycolytic gene expression in embryos that are cultured in media that includes 10 mM glucose and 1 mM α-KB, but lacks pyruvate and lactate. **(K-K”)** The levels of GAPDH **(K, K’)** increase following pyruvate withdrawal. **(K”)** Quantitation of **(K, K’)**. **(L-L”)** The levels of GLUD1 **(L, L’)** do not increase following pyruvate withdrawal. **(L”)** Quantitation of **(L, L’)**.

**Figure S6, related to Figure 7:**
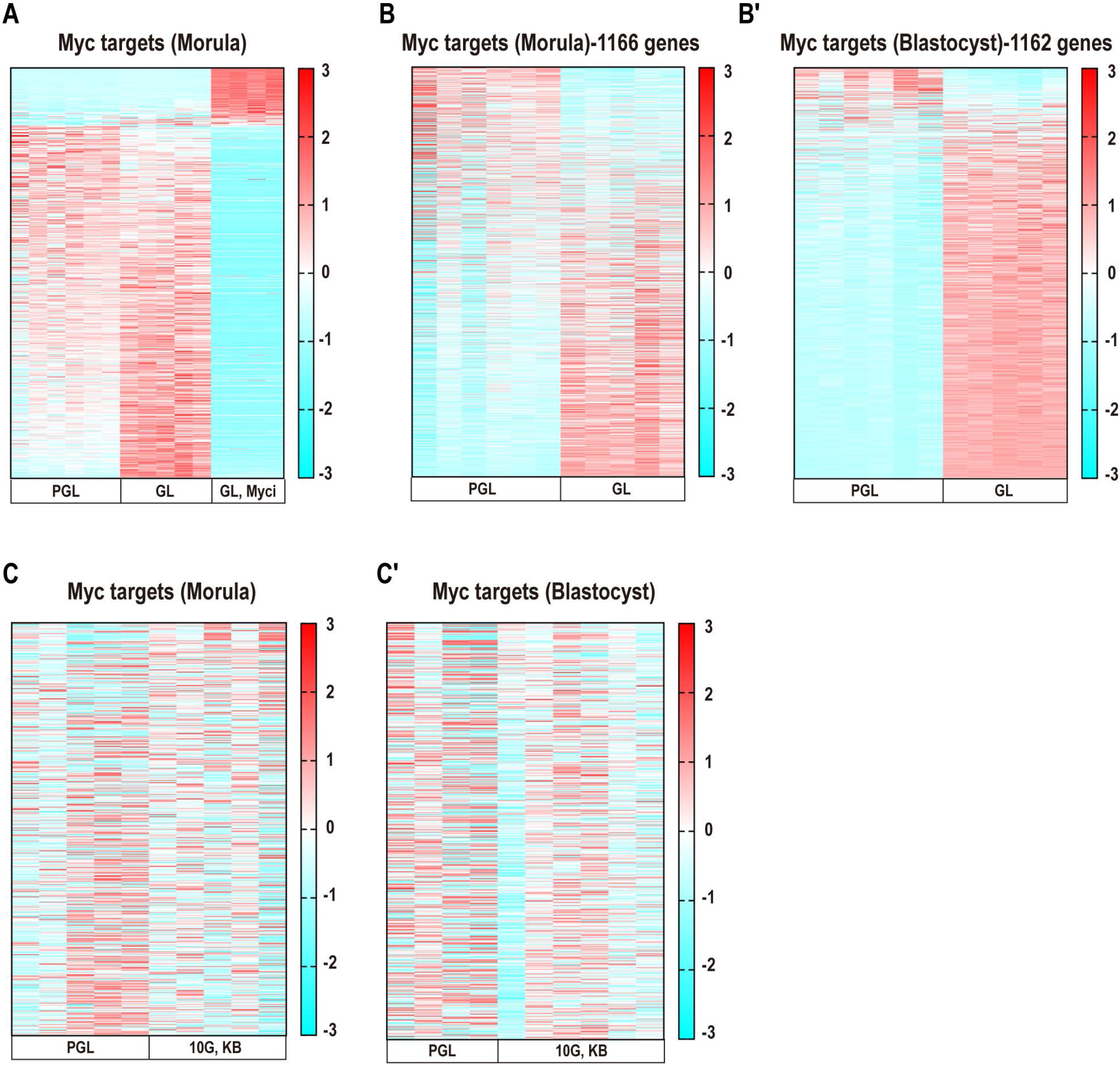
Mechanisms of metabolic reprogramming during reductive stress. **(A)** In the presence of the Myc inhibitor (20nM KJ-Pyr-9) a majority of the Myc targets decrease. **(B, B’)** A majority of Myc targets (total = 1166 genes, list 2) increase in expression at the morula stage **(B)** and more so in blastocysts **(B’)** following pyruvate withdrawal at the late 2C stage (50h). **(C, C’)** Myc targets do not increase in expression in embryos cultured from the late 2C stage in a medium that contains 10mM glucose and 1mM α-KB, but lacks in pyruvate and lactate until the morula **(C)** and blastocyst **(C’)** stage. The Myc targets used are the same targets that are used in Figure 7J, J’ (list 1).

